# Genomic prediction models based on a large-scale recombinant population allow rapid breeding of desired genotypes

**DOI:** 10.1101/2025.09.14.676066

**Authors:** Toshiyuki Sakai, Hiroki Takagi, Tomoaki Fujioka, Yuuki Oota, Shinsuke Nakajo, Hiroki Yaegashi, Kaori Oikawa, Hiroe Utsushi, Kazue Ito, Satoshi Natsume, Motoki Shimizu, Takumi Takeda, Ryohei Terauchi, Akira Abe

**Author notes:** These authors contributed equally to this work (Toshiyuki Sakai, Akira Abe). Ishikawa Prefectural University, Nonoichi, Ishikawa 921-8836, Japan (Hiroki Takagi). Iwate Prefectural Agricultural College, Kanegasaki, Iwate 029-4501, Japan (Tomoaki Fujioka). Agriculture, Forestry and Fisheries Department, Iwate Prefectural Government, Morioka, Iwate 029-4501, Japan (Yuuki Oota). (T.S.); (R.T.), (A.A.).

## Abstract

Rapid development of cultivars with optimal genome-wide allele combinations is essential for addressing agricultural challenges. However, constructing desired genotypes through conventional cross-breeding requires many generations, calling for a more efficient methodology. Here, we present a rapid breeding strategy that combines a large-scale recombinant population as starting material with interpretable genomic prediction models trained on that population. This approach enables efficient construction of target genotypes optimized for multiple traits in cultivars. To validate our strategy in rice (*Oryza sativa*), we established a nested association mapping population of 2,787 recombinant inbred lines derived from an elite cultivar ’Hitomebore’ and 19 diverse donors. We built highly accurate genomic prediction models using this population. We then used the models to estimate haplotype-specific effects and account for trade-offs among yield-related traits, and selected optimal lines and designed breeding schemes. Crossbreeding based on these schemes produced rice lines with the target genotypes for multiple yield-related traits, supporting the predicted effects and validating the effectiveness of our strategy. This genomic breeding approach provides a general framework for rapidly breeding cultivars able to meet the challenges posed by a changing environment.

## INTRODUCTION

Meeting global food demand under a changing environment requires flexible and efficient breeding strategies that are capable of producing new cultivars of major food crops with desired traits (*1–4*). One route to producing such cultivars is to pyramid favorable alleles for economically important traits, scattered across the genome, onto the genomic background of an elite cultivar (*5–8*). Because breeding objectives shift with climate change, sustainability concerns, and consumer preferences, the desired genotype itself changes accordingly (*7*). However, conventional breeding approaches require many generations of crossing and selection to reach a target genotype and often struggle to meet these diverse and shifting demands. More efficient approaches are therefore needed that can be readily implemented in breeding programs to rapidly produce new cultivars with genotypes optimized for multiple traits.

Constructing such genotypes requires accurately estimating the genetic effects of genomic regions on the traits of interest, selecting favorable haplotypes for introgression while accounting for genetic trade-offs among traits, and introducing the target haplotypes through cross-breeding (*7*, *9*). Use of a nested association mapping (NAM) population can accelerate all these steps. NAM populations consist of many recombinant inbred lines (RILs) derived from crosses between genetically diverse varieties and a common parental line, which allows quantitative trait loci (QTLs) and genes underlying trait variation to be identified (*10*, *11*). NAM populations are also well suited for building accurate genomic prediction models that capture genetic effects on trait values at high resolution, enabling the prediction of optimal genotypes across traits (*7*, *12*). Furthermore, once established, the inbred lines themselves serve as permanent pre-breeding material, ready to rapidly produce the target genotypes on demand (*13*). Combining NAM populations with genomic prediction models thus offers an efficient route to constructing such genotypes.

In this study, we apply genomic prediction to a NAM population for rapid breeding of cultivars with target genotypes. We established a rice NAM population and used it to develop interpretable genomic prediction models, which we then applied to practical breeding programs to efficiently construct cultivars with target genotypes optimized for multiple traits (Fig. 1).

**Fig. 1.**
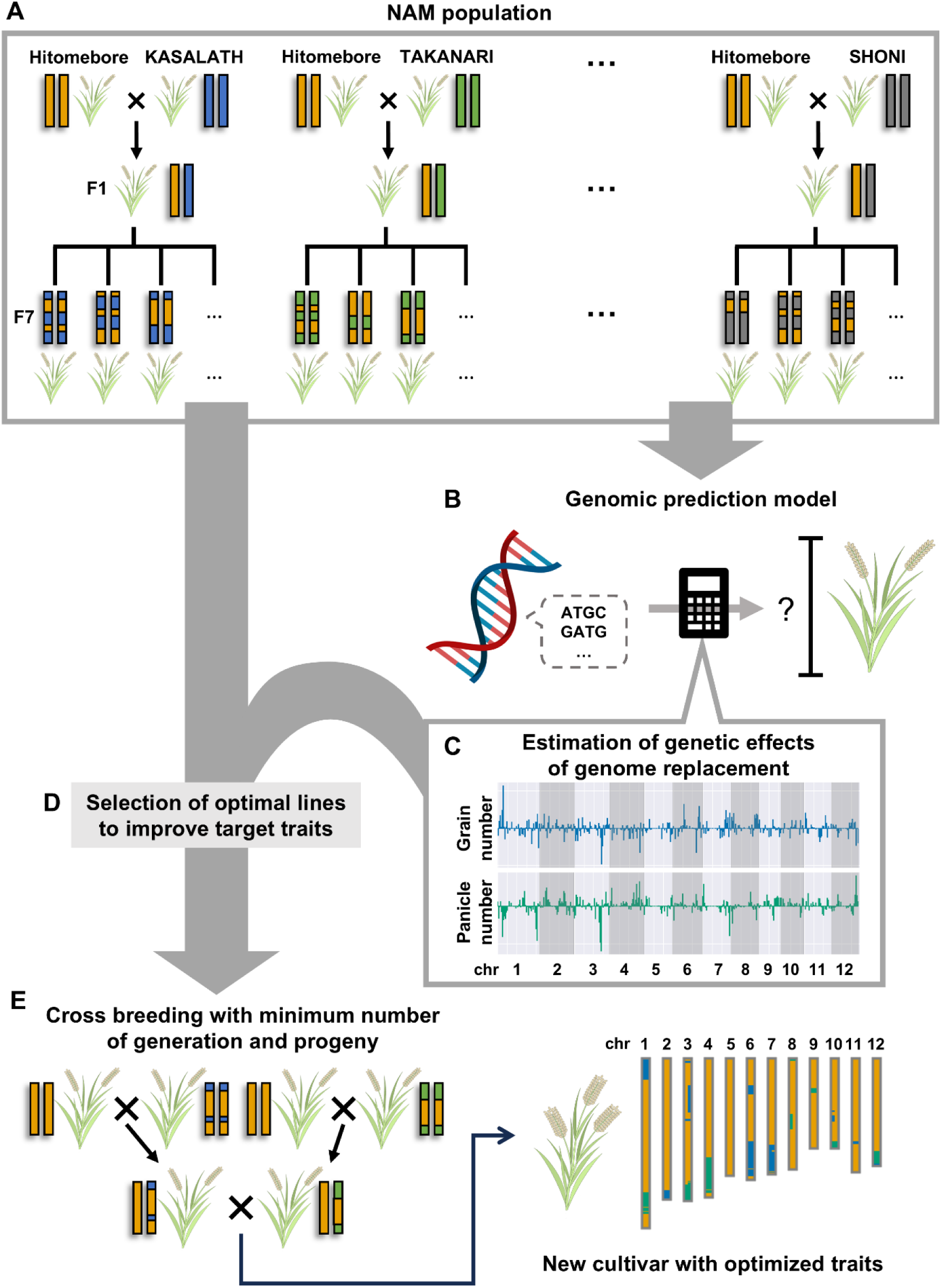
Flowchart of the breeding strategy combining interpretable genomic predictions and crossbreeding of selected recombinant inbred lines from a nested association mapping population. (A) A nested association mapping (NAM) population is generated using the local elite cultivar Hitomebore as the common parent. (B) Interpretable genomic prediction models are constructed based on the phenotype–genotype relationships detected in the NAM population. (C) The genetic effects caused by the replacement of a given genomic region from the common parent by the equivalent region from the donor parents are estimated by the genomic prediction models. (D) Optimal recombinant inbred lines (RILs) are selected from the NAM population based on their estimated genetic effects. (E) The selected RILs are used for rapid breeding to produce a new cultivar with the desired phenotypes for multiple traits.

## RESULTS

### Use of a rice NAM population for genomic prediction and breeding

We established a NAM population in rice consisting of 19 RIL populations for a total of 2,787 lines. The RILs were derived from crosses between the *temperate japonica* cultivar ‘Hitomebore’, an elite cultivar from Northern Japan, as the common parent, and 19 genetically diverse donor cultivars representing the major rice phylogenetic clades (*aus*, *indica*, *temperate japonica*, *tropical japonica*) (Fig. S1). We grew the entire NAM population in the experimental field of Iwate Agricultural Research Center, Iwate, Japan, and recorded the phenotypic values for five traits over 2–6 years: grain number per panicle (hereafter abbreviated as grain number), panicle number, heading date, leaf length, and leaf width. Grain size was measured one year. The distribution of phenotypic values in each RIL population exhibited transgressive segregation for all six traits measured (Fig. S2). This pattern indicates that the parental lines have alleles with positive and negative effects on the trait values, which can surpass those of the parents depending on the genomic composition of the individual line.

We also investigated the potential for genetic trade-off relationships among the average phenotypic values for yield-related traits (grain number, panicle number, and grain size) over the years across the entire NAM population grown in an identical environment. Plants producing more grains per panicle tended to carry fewer panicles and had smaller grains, suggesting the presence of genetic trade-off among these three agronomic traits (Fig. 2). Therefore, the genotype with a higher total grain mass per plant (estimated as panicle number × grain number × grain size) should be searched at the Pareto optimality front of the phenotypic values for the three constituent traits (Fig. 2). Also, RILs with extreme phenotypic values for an individual trait are not suitable for conventional cultivation. Thus, multiple traits need to be controlled simultaneously in an optimized balance to maximize total yield.

**Fig. 2.**
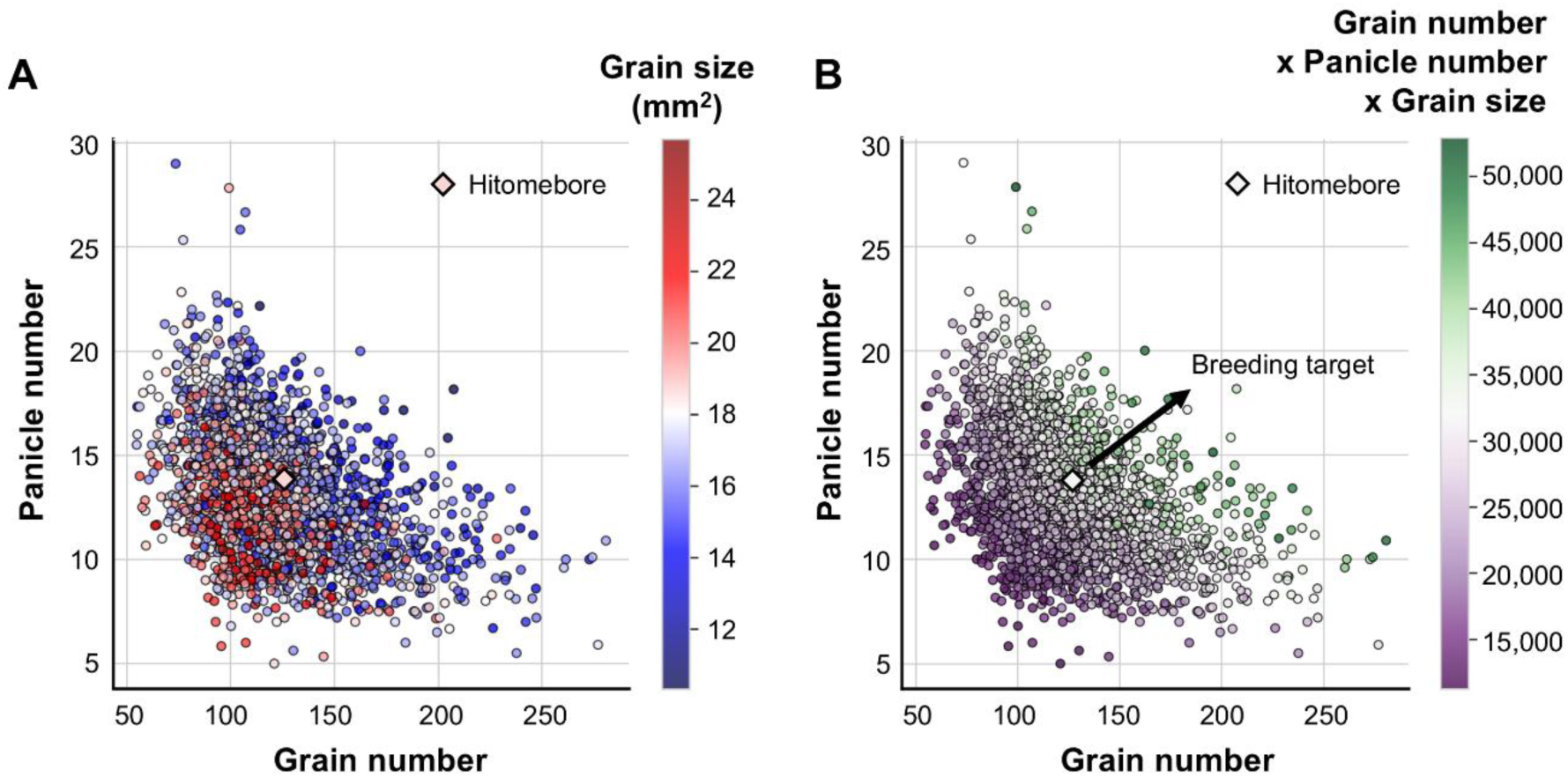
Trade-off relationships among the values for three agronomically important traits across 2,787 RILs. Grain size, grain number, and panicle number were measured for 2,787 RILs of the NAM population developed using Hitomebore as the common parent. (**A**) Scatterplot showing the relationships among grain number (*x*-axis), panicle number (*y*-axis), and grain size (color-coded) in the NAM population. (**B**) Scatterplot showing the relationships among grain number (*x*-axis), panicle number (*y*-axis), and yield (approximated by calculating grain number × panicle number × grain size; color-coded). The black arrow indicates the vector of changes in trait value that meet our breeding target, which is to improve grain number and panicle number simultaneously without allowing them taking extreme phenotypic values in individual traits.

To obtain the genotype information of all RILs, we resequenced their whole genomes and identified 2,267,856 single-nucleotide polymorphisms (SNPs) that aggregated into 57,085 haplotype blocks (Fig. S1). To infer the genotypic diversity of the NAM population, we performed a principal component analysis (PCA) based on the genotypes at these haplotype blocks. RILs derived from a *temperate japonica* cultivar showed very low genetic and phenotypic diversity, as illustrated by their tight clustering in the PCA plots (Fig. S3). RILs derived from *indica* donor cultivars or *tropical japonica* cultivars exhibited a lower genetic diversity than those derived from an *aus* cultivar. However, phenotypic diversity for the agronomic traits grain number and panicle number was equally large in the three groups (Fig. S3). The inconsistency between the genetic and phenotypic diversity observed for RILs derived from *indica* and *tropical japonica* cultivars suggests that their trait values are determined by a relatively small number of genomic regions with strong effects. We consider that the NAM population developed for this study contains sufficient genetic diversity to improve the agronomic traits of the Hitomebore cultivar, the common parent of the NAM population, as demonstrated below.

### Accurate genomic prediction models based on the NAM population

With the phenotypic and genotypic data from the 2,787 RILs of the rice NAM population in hand, we built genomic prediction models for improving the rice cultivars. To build suitable models, we tested eight statistical models, namely Lasso, Elastic Net, Genomic Best Linear Unbiased Prediction (GBLUP), Random Forest, Light Gradient Boosting Machine (LightGBM), BayesA, BayesB, and BayesC. We evaluated the accuracy of the models by a five-fold cross-validation, based on 20% of data from the NAM population as a test set (Fig. 3A). There were no significant differences among the statistical models except for Random Forest, which clearly performed worse than the other seven models (Fig. S4). The models built by Elastic Net showed high accuracy for all traits, with high Pearson’s correlation coefficients (*r*) between the measured and predicted phenotypic values: *r* = 0.84 for grain number, *r* = 0.86 for panicle number, *r* = 0.85 for grain size, *r* = 0.88 for leaf width, *r* = 0.76 for leaf length, and *r* = 0.83 for heading date (Fig. 3B, Fig. S5). For all traits, genomic prediction models using fewer than 1,000 haplotype blocks with high contributions to phenotypes attained nearly the same accuracy as those using all haplotype blocks (Fig. S6). We thus built the models only with haplotype blocks most relevant to each trait to enhance interpretability. Hence, we employed the Elastic Net model using haplotype blocks selected from the entire NAM dataset to build our model for subsequent analysis.

**Fig. 3.**
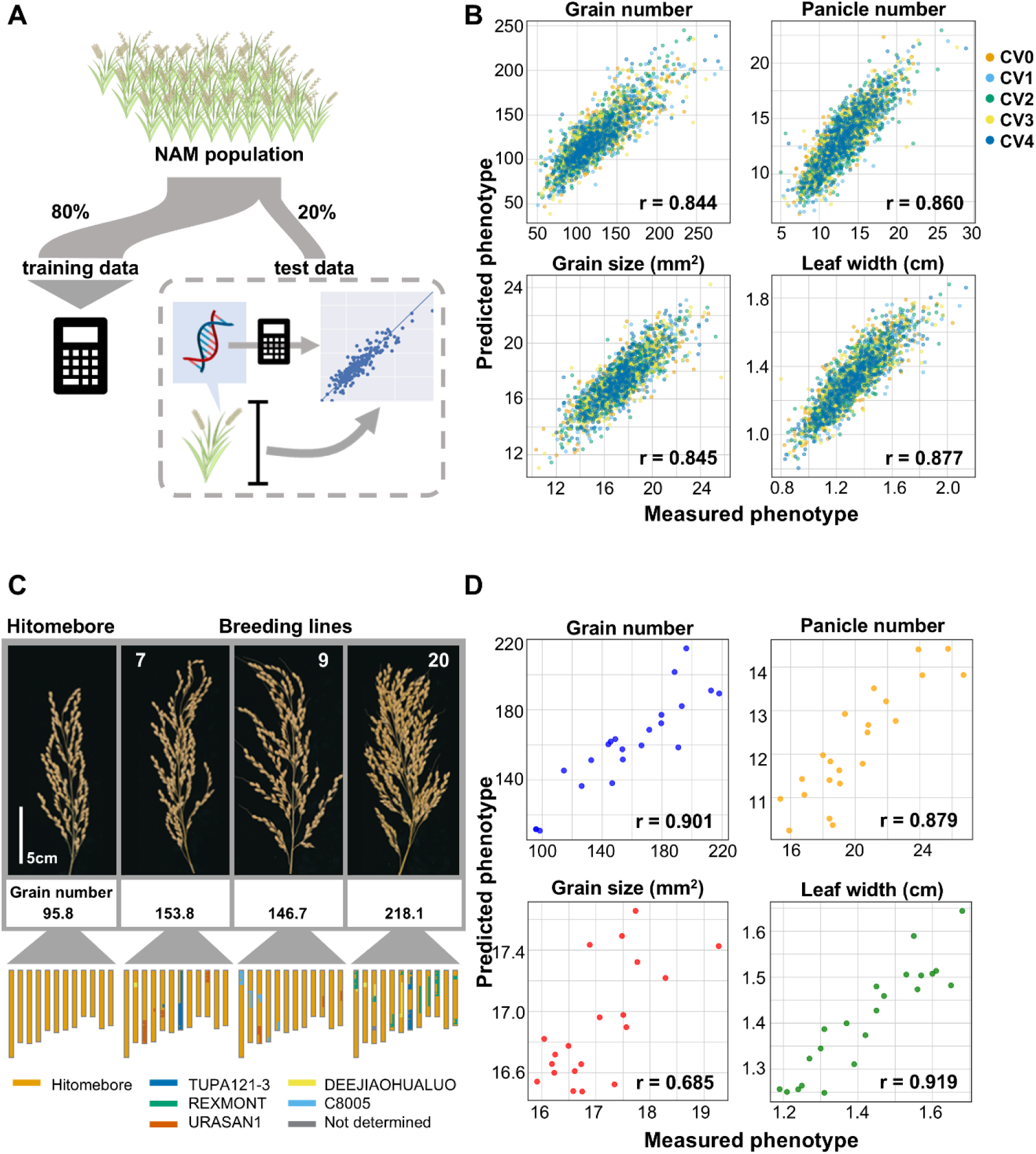
Genomic prediction models constructed using data collected from the NAM population allow accurate predictions of agronomically important traits. (**A**) Diagram of the construction of genomic prediction models (training data) and the evaluation of their prediction accuracy (test data). (**B**) Scatterplots showing the relationships between the measured and predicted phenotypic values in the test data for the four traits: grain number, panicle number, grain size, and leaf width, as predicted by Elastic Net. The mean Pearson’s correlation coefficients (*r*) between the measured and predicted phenotypic values from five-fold cross-validation (CVs) are shown to indicate the prediction accuracy. (**C**) Representative photographs of panicles and the graphical genotypes of Hitomebore and three representative breeding lines harboring genomic fragments from more than two donor cultivars. (**D**) Scatterplots showing the relationships between measured and predicted phenotypic values for the 21 breeding lines carrying genomic fragments from more than two donor lines for the four traits.

Each RIL used for genomic prediction modeling carries genomic fragments derived from only two cultivars: Hitomebore and the donor cultivar. The accuracy of such predictions when applied to more complicated genotypes with genomic fragments originating from more than two cultivars is unknown. To evaluate the robustness of the models against such genotypes, we selected 21 new breeding lines that contain genomic fragments derived from several of the same founders used for developing the NAM population (Fig. 3C, Fig. S7). When applied to the phenotypic values of grain number, panicle number, grain size, and leaf width, our genomic prediction models maintained high accuracy for grain number (*r* = 0.90), panicle number (*r* = 0.88), and leaf width (*r* = 0.92), while being less accurate for grain size (*r* = 0.69) (Fig. 3D). These results indicate that the genomic prediction models developed here based on bi-parental RILs are robust enough to also predict phenotypes from lines with more complex genome configurations.

To further investigate the versatility of our approach developed for a rice NAM population, we applied the Elastic Net model based on all haplotype blocks to genomic prediction using NAM datasets from maize (*Zea mays*), soybean (*Glycine max*), and sorghum (*Sorghum bicolor*) for which genotypes and phenotypic values are publicly available (*10*, *14*, *15*). The genomic prediction models applied to 19 traits consistently showed high Pearson’s correlation coefficient values indicative of high predictive accuracy (*r*): 8 of the 10 traits in maize, all 7 traits in soybean, and the 2 traits in sorghum showed *r* > 0.7 (Fig. S8). We conclude that NAM populations generally support building highly accurate genomic prediction models to estimate phenotypic values.

### Interpretation of the models and developing an optimal breeding plan for target genotypes to improve multiple traits

Using the genomic prediction models and the RILs, we developed a pipeline for rapid breeding of target genotypes by crossbreeding, in three steps: (1) identification of genomic regions positively or negatively affecting the target trait and overall yield, (2) selection of suitable RILs for crossbreeding with genomic regions positively affecting the trait, and (3) rapid breeding over few generations and with small population sizes (Fig. 1B-E).

As a proof of concept, we applied the above pipeline to improve Hitomebore, the common parent used for the NAM population, for grain number and panicle number simultaneously using two donor lines. We first estimated the effects of all genomic regions on three yield-related traits, namely grain number, panicle number, and grain size, by interpreting the genomic prediction model. Specifically, we estimated the effect of replacing a haplotype block from Hitomebore with that from a donor cultivar, based on regression coefficients of the model (Fig. 4A). Owing to the trade-off among agronomic traits, some genomic regions have positive effects on one agronomic trait but negative effects on other yield-related traits. Such genomic regions from the donor parent may be detrimental for total yield, even though they improve a single trait. To estimate the effect of replacing a genomic region on overall yield, we calculated the sum of Z-scores for all effects on grain number, panicle number, and grain size as the provisional effect on yield (Fig. 4A). From these estimated effects, we assessed which genomic regions from which donor should be introduced into the Hitomebore genome to improve the target traits without negatively affecting total yield (Fig. 4A).

**Fig. 4.**
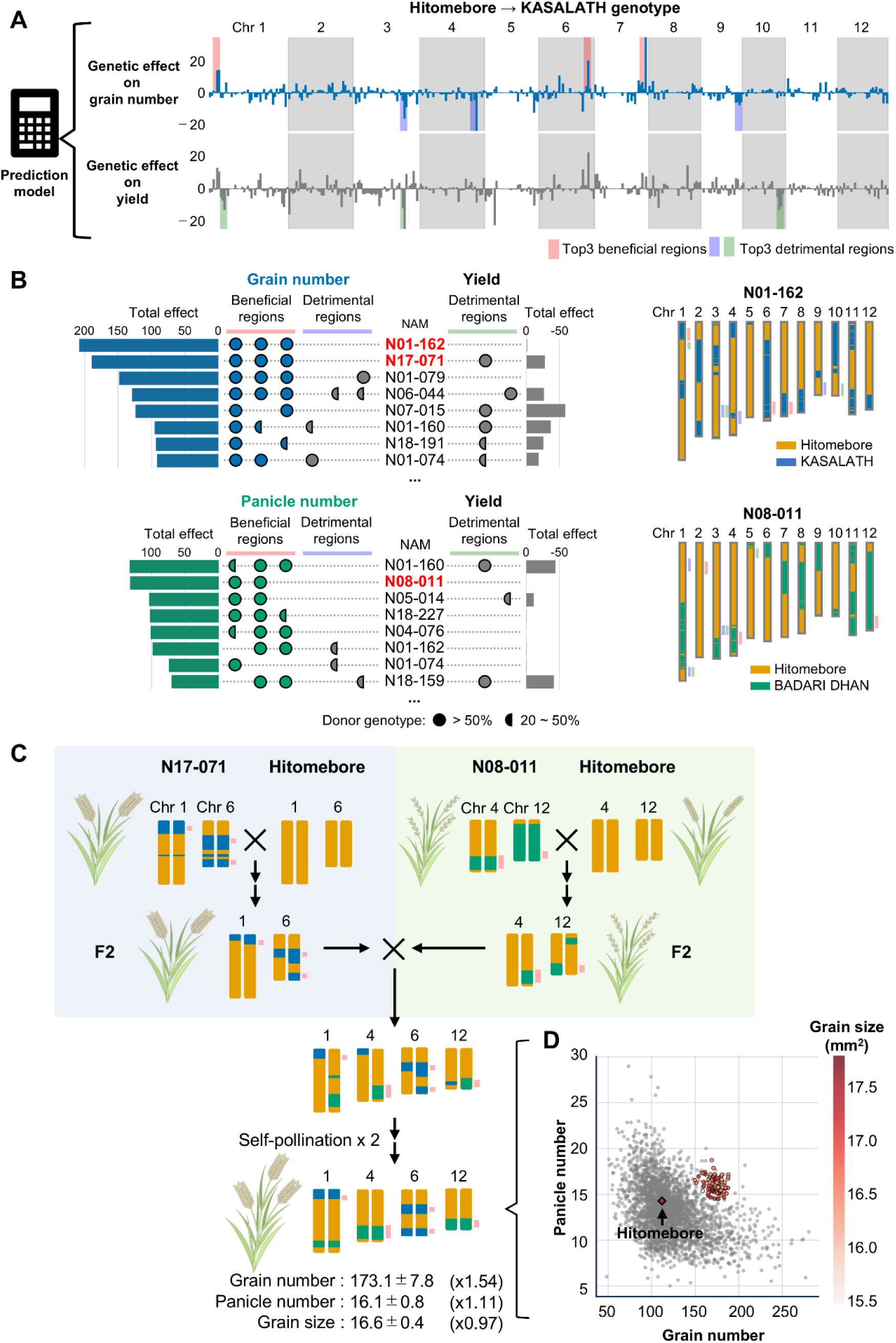
Overview of our breeding strategy. Interpretable genomic prediction models select the best RILs as the starting materials for breeding; these RILs are then used to rapidly crossbreed the target cultivar, which is optimized for multiple traits. **(A)** An example of estimated genetic effects for grain number and yield when the genomic region from Hitomebore is replaced with the equivalent region from KASALATH. The top three genomic regions positively contributing to the target trait (beneficial regions) and the top three genomic regions negatively affecting the trait and total yield (detrimental regions) were selected based on their estimated effects, as indicated by the genomic prediction models. **(B)** The optimal RILs that improve grain number and panicle number are selected based on the total genetic effects of all beneficial and detrimental genomic regions, with RILs containing the highest number of beneficial genomic regions and the lowest number of detrimental genomic regions (or none) chosen for crossbreeding. Graphical genome representations of two exemplar optimal RILs are shown to the right. **(C)** Diagram illustrating a breeding plan using two RILs, N17-071 and N08-011. In the breeding plan, each RIL is crossed to Hitomebore, followed by a cross between individuals from the two resulting F2 populations, and two rounds of self-pollination to generate a line homozygous for the selected regions **(D)** Scatterplot showing the predicted phenotypic values for grain number (*x*-axis) and panicle number (*y*-axis) in the target line obtained from 100 simulations (red circles) and the measured phenotypic values for the RILs in the NAM population, indicated as gray circles. The predicted phenotypic values for Hitomebore are indicated by the arrowhead.

Based on the information of genomic regions related to the trait variation and the total yield, we selected one RIL suitable for raising grain number and another RIL suitable for improving panicle number (Fig. 4B). We selected RILs harboring three genomic regions with the highest total effects from the donor parent and at the same time lacking the three genomic regions with the strongest negative effects on the target trait and total yield (Fig. 4B). For this selection step, we also prioritized RILs with a higher proportion of genomic fragments derived from the Hitomebore parent to retain most genomic regions as close to Hitomebore as possible (Fig. S9).

To improve grain number, we selected RIL N01-162, whose genotypic composition had the largest total positive effect on grain number. We also selected RIL N17-071, which had the second-largest total positive effect on grain number in the targeted genomic regions (Fig. 4B). To improve panicle number, we selected RIL N08-011, which had the largest positive effects on panicle number conferred by two beneficial genomic regions (Fig. 4B). The genome of each of the three RILs was over 65% derived from Hitomebore (Fig. S9) and also carried the Hitomebore genotype at most of the genomic regions whose equivalent regions in the donor genomes are associated with detrimental yield effects (Fig. 4B). Using the selected RILs, we designed a breeding scheme involving N17-071 and N08-011 (Fig. 4C), and another one involving N01-162 and N08-011 (Fig. S10).

We considered a breeding plan to rapidly accumulate beneficial genomic regions for grain number and panicle number into Hitomebore from the two selected RILs for each scheme. The breeding scheme was as follows: (1) cross the RIL selected for improving grain number to Hitomebore and generate an F2 population (named grain number-F2) with a greater proportion of the Hitomebore genome than the starting RIL; do the same with the RIL selected for panicle number (named panicle number-F2); (2) cross a suitable individual from grain number-F2 to a suitable individual from panicle number-F2 to combine their respective beneficial genomic regions; and (3) propagate by self-pollination to generate plants homozygous at most genomic locations (Fig. 4C, Fig. S10).

To evaluate whether the target lines could be produced with a minimal number of individuals and generations, we simulated the breeding program. For this purpose, we modeled the number and positions of recombination events per cross based on the actual recombination patterns observed in the NAM population (Fig. S11). We designed ∼50 markers that distinguish between the donor and Hitomebore genomes and used them to select of individuals to be used in the subsequent cross (Fig. S12). Based on the simulation, we estimated that the target genotype may be obtained in two years with the rice speed-breeding scheme, which allows for three generations per year (Fig. S10) (*16*). Our simulation also indicated that fewer than 100 plants per cross, the maximum number attained in speed breeding, are sufficient to bring the five donor genomic regions into the Hitomebore genome (Fig. S10).

Our prediction showed that the target line produced using RILs N17-071 and N08-011 would produce 54% more grains and 11% more panicles than Hitomebore, with grains about the same size (97%) as those from Hitomebore. This target line would contain genomic regions covering over 84.2% of the Hitomebore genome (Fig 4C, Fig. S10). Importantly, the predicted target line had phenotypic values close to the Pareto optimality front for the panicle number–grain number trade-off (Fig. 4D). Using RILs N01-162 and N08-011, we predicted another target line with similar phenotypes, achieving 47% more grains, 9% more panicles, and grains 95% the size of those from Hitomebore, with a genome composition encompassing over 83.1% of genomic regions derived from Hitomebore (Fig. S10). The above genomic prediction pipeline applied to RILs from the NAM population suggests the possibility of rapid construction of target genotypes improving grain number and panicle number without significant negative trade-off effect while retaining a high proportion of genomic regions derived from Hitomebore.

### Experimental validation of the breeding plan

To validate the rapid breeding pipeline, we selected 10 F2 individuals each from the crosses N17-071 × N08-011 and N01-162 × N08-011. These F2 individuals inherit variable numbers of the five target regions from the RILs judged by the whole-genome sequencing data. We obtained the F3 lines derived from the self-pollination of each selected F2 individual and evaluated their phenotypes to investigate the effects of the five beneficial genomic regions (Fig. 5, Fig. S13). We used the average phenotypic values of all individuals from each F3 line as representative values for their corresponding F2 parent. The measured phenotypic values among the F2 individuals derived from N17-071 × N08-011, as well as those from the parental RILs, showed a high correlation with the values predicted from the genotypes for grain number (*r* = 0.85), panicle number (*r* = 0.69), and grain size (*r* = 0.65) (Fig. S14). The phenotypic values for the F2 individuals were clearly higher than those for Hitomebore and reflected the effects of beneficial genomic regions (Fig. 5). F2 plants carrying all five beneficial genomic regions showed higher grain number and panicle number than Hitomebore, with grain size remaining ∼90% that of Hitomebore (Fig. 5). The 10 F2 individuals derived from N01-162 × N08-011 also showed clear effects from the accumulation of beneficial genomic regions and high correlations between measured and predicted phenotypic values for grain number (*r* = 0.96), panicle number (*r* = 0.79), and grain size (*r* = 0.64) (Figs. S13 and S14). These results demonstrate that superior cultivars can be produced by optimizing multiple traits based on genomic information and by rapid breeding for combining genotypes.

**Fig. 5.**
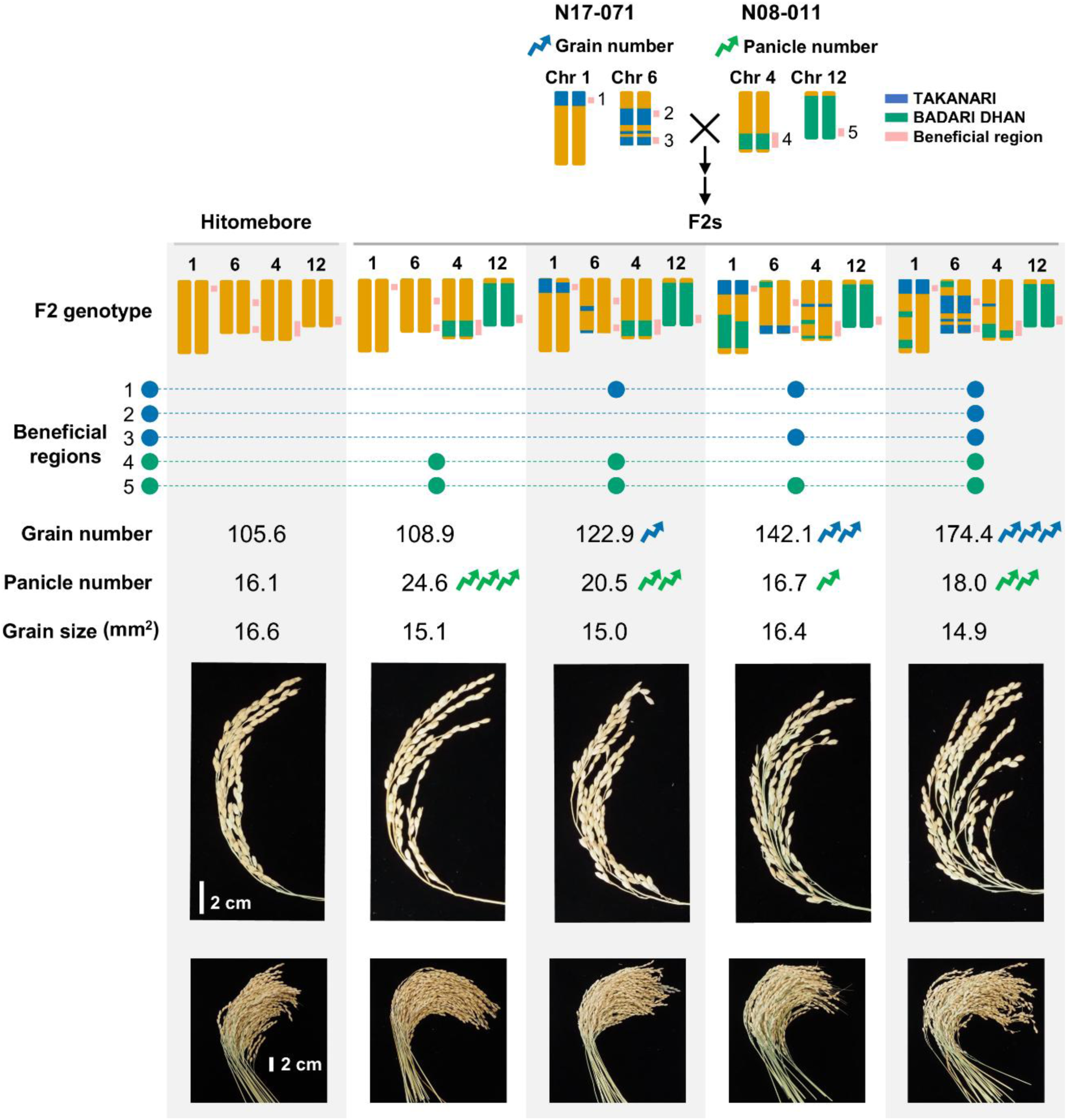
Validation of the phenotypes in lines optimized by genomic prediction. Diagram illustrating the generation of the F2 population from a cross between the optimal RILs N17-071 and N08-011. Graphical representations of their genotypes at the target genomic regions on the respective chromosomes, the phenotypic values for grain number, panicle number, and grain size, and images of panicles from representative F2 plants are shown. The phenotypic values for each F2 plant represent the average values from their F3 progeny.

## DISCUSSION

We developed an efficient genomic breeding strategy based on interpretable genomic prediction models and a NAM population that enabled the construction of an optimal breeding plan to rapidly generate new cultivars with desired genotypes (Fig. 1). This strategy differs from conventional breeding approaches that focus on continuous genetic gain through long-term selection or on the improvement of a single trait based on markers without considering the effect size of QTL replacement and the trade-offs among multiple traits (*17*). Our strategy provides a clear plan to develop target genotypes that comprehensively optimize multiple traits in new cultivars over a short timeframe, provided that a NAM population has already been established for that species. In addition, we built highly accurate genomic prediction models for maize, soybean, and sorghum, using the same method, confirming that our strategy is generally applicable to crop improvement. Establishing a NAM population using a local elite cultivar as a common parent enables rapid genetic improvement of that cultivar through our pipeline, thereby enhancing agricultural productivity. Moreover, utilizing genetically diverse resources, such as core collections and pangenomes established in recent years, to inform the generation of NAM populations should expand the potential for genetic improvement achievable through this approach. Leveraging crop genetic diversity and state-of-the-art modeling, we believe our strategy helps achieve high crop productivity in a rapidly changing global environment.

## MATERIALS AND METHODS

### Plant materials and growth conditions

To establish a nested association mapping (NAM) population, the *japonica* rice (*Oryza sativa*) cultivar ‘Hitomebore’ was used as the common parent and was crossed to 19 varieties: eight *aus* varieties (‘KASALATH’, ‘SHONI’, ‘TUPA 121-3’, ‘SURJAMUKHI’, ‘RATUL’, ‘BADARI DHAN’, ‘KALUHEENATI’, and ‘C8005’), three *indica* varieties (‘KEIBOBA’, ‘DEEJIAOHUALUO’, and ‘TAKANARI’), six *tropical japonica* varieties (‘JAGUARY’, ‘REXMONT’, ‘URASAN 1’, ‘TUPA 729’, ‘SESIA’, and ‘NERICA1’), and two *temperate japonica* varieties (‘MOUKOTO’ and ‘NORTAI’). These varieties were obtained from the National Agriculture and Food Research Organization World Rice Core Collection (*18*) or our own collection. For each cross, the resulting F1 plant was self-pollinated to obtain the F2 generation, which was used to generate the F7–F9 generations by the single-seed-descent method. The code names and number of lines for each recombinant inbred line (RIL) are summarized in Data S1. These RIL populations were planted in experimental paddy fields at the Iwate Agricultural Research Center in Kitakami, Iwate Prefecture, Japan (latitude 39°35′N, longitude 141°11′E, altitude 89 m), in 2013–2019 for phenotyping. Fifteen seeds per RIL, together with their parental lines, were germinated in water and sown in 2 × 2-cm plug trays filled with synthetic nursery soil (Gosei-Baido L, Sanken-soil Co. Ltd., Japan). At 25–30 days after sowing, seedlings from each RIL and parental line were transplanted into paddy fields (one seedling per hill) at a density of 22.2 hills per m^2^.

To validate the genomic prediction model, we used breeding lines derived from Hitomebore with introgressed genomic segments from donor varieties, including several of the NAM founders (e.g. TUPA 121-3, DEEJIAOHUALUO, C8005, URASAN 1, REXMONT) as well as additional donors such as TOYAMA 73, and DUNGHAN SHALI. These segments were introduced through multiple rounds of crossing, including backcrossing to Hitomebore. Some lines carried segments from a single donor, while other lines contained pyramided fragments from multiple donors. The combinations of donor varieties used are listed in table S1. The lines were transplanted in 2020 into a paddy field (three seedlings per hill) at a density of 22.2 hills per m^2^. For the validation of the breeding plan, F2 populations were generated from two crosses: N17-071 × N08-011 and N01-162 × N08-011. 10 F2 plants were selected based on the genotypes of beneficial genomic regions. From each F2 plant, 7–49 F3 progeny were generated and transplanted into a paddy field (one seedling per hill) at a density of 22.2 hills per m^2^ in 2024. Basal fertilizer was applied at a rate of 30 kg nitrogen (N), 30 kg phosphorus (P_2_O_5_), and 30 kg potassium (K_2_O) per hectare (2013–2019), or 60 kg ha⁻¹ each of N, P₂O₅, and K₂O for the 2020 and 2024 experiments.

### Phenotyping

Values for the following six traits were measured for all RILs and the parental lines: leaf length (cm), leaf width (cm), grain number, panicle number, grain size (mm^2^), and heading date. Leaf length and leaf width were measured using the flag leaf of the main culm in 2013–2016. Grain number was counted from a panicle on the main culm in 2016 and 2017. Panicle number per plant was counted in 2014, 2016, and 2017. Grain size as measured in 2016. Heading date was counted as days to heading after sowing in 2013–2017. The day when 50% of the individuals in a RIL had visible panicles was defined as the heading date of that line. The values for leaf width, leaf length, grain number, panicle number, and heading date were also collected for the lines in RIL populations N05, N11, N12, N17, and N18 in 2018 and 2019. For all phenotypic data, six individuals per RIL were assessed; their mean values were calculated for each year. For subsequent analyses, the averages across all investigated years were used for all traits. For breeding lines and F3 progeny used for validation, grain number, panicle number, and grain size were measured using the same method as above, except that panicle number for breeding lines was counted per hill.

### Genotyping analysis

To obtain the genotypes of the RILs, whole-genome resequencing of the parental lines and all RILs was performed. First, genomic DNA was extracted from fresh leaves using an Agencourt Chloropure Kit (Beckman Coulter, California, USA). Then, DNA concentration was quantified using Invitrogen Quant-iT PicoGreen dsDNA Assay Kits (Thermo Fisher Scientific, Massachusetts, USA). For the parental lines, library construction was performed using a TruSeq DNA PCR-Free Library Prep Kit (Illumina, California, USA). These libraries were sequenced on three Illumina platforms: MiSeq (75-bp paired-end reads), HiSeq(150-bp paired-end reads), and NextSeq500 (250-bp or 300-bp paired-end reads). For all RILs, library construction was performed using house-made sequencing adapters and indices. These libraries were sequenced on an Illumina NextSeq500 instrument as 75-bp paired-end reads. First, adapter sequences were removed using FaQCs v2.08 (*19*). Then, PRINSEQ lite v0.20.4 (*20*) was used to remove low-quality bases with the option “-trim_left 5 -trim_qual_right 20 -min_qual_mean 20 -min_len 50”. In addition, 250-bp or 300-bp reads were trimmed to 200 bp by adding the option “-trim_to_len 200”.

Genotyping was conducted separately for each RIL population derived from different parental combinations, using the single-nucleotide polymorphism (SNP) set specific to each pair of parental lines. The quality-trimmed short reads were aligned against the reference genome using Burrows-Wheeler Aligner (BWA) v0.7.17 (*21*). Os-Nipponbare-Reference-IRGSP-1.0 was used as the reference genome (*22*). After mapping, the BAM files were sorted and indexed using SAMtools v1.10 (*23*), which were subsequently subjected to variant calling. Variant calling for the parental lines was performed according to the “GATK Best Practices for Germline short variant discovery” pipeline (*24*) (https://gatk.broadinstitute.org/), which includes a base quality score recalibration (BQSR) step, two variant calling steps with HaplotypeCaller in GVCF mode and GenotypeGVCFs commands, and a filter variant step with VariantFiltration command with the option “QD < 2.0 || FS > 60.0 || MQ < 40.0 || MQRankSum < -12.5 || ReadPosRankSum < -8.0 || SOR > 4.0”. From the resulting VCF file, only biallelic SNPs were retained when (i) both parental lines had homozygous alleles, (ii) the genotypes of the two parental lines were different, and (iii) both parental lines had a read depth (DP) of at least 8. The positions of selected SNPs were converted to a BED file (position.bed) using the query command in BCFtools v1.10.2 (*25*). For SNP genotyping of the RILs, the VCF files were generated as follows: (i) BCFtools mpileup command with the options “-t DP,AD,SP -A -B -Q 18 -C 50 -uv -l position.bed”; (ii) BCFtools call command with the option “-P 0 -A -c -f GQ”; (iii) BCFtools filter command with the option “-v snps -i ‘INFO/MQ> = 0 & INFO/MQ0F< = 1 & AVG(GQ)> = 0’”; and (iv) BCFtools norm command with the option “-m+both”. The resulting variants were imputed based on the parental genotypes using LB-impute (*26*). Finally, the imputed genotypes from all 20 RIL populations were integrated into a unified genotype matrix for downstream analysis. In total, a total of 2,267,856 SNPs were identified. The number of SNPs between the common parent, Hitomebore, and each donor parent is summarized in Data S1. To identify the number of recombination events, the number of positions where the genotypes of a genomic region spanning more than 500 kb switched from the Hitomebore genotype to the donor genotype was counted. The average number of recombination events in the whole genome was 27.1. The number of recombination events for each RIL population is also summarized in Data S1.

To validate the genomic prediction model and breeding plan, whole-genome resequencing was performed on the 21 breeding lines and two F2 populations derived from crosses between three RILs. Genomic DNA was extracted from fresh leaves using a NucleoSpin Plant II kit (Macherey-Nagel, Düren, Germany). Libraries were prepared using a Collibri ES DNA Library Prep Kit for Illumina system (Thermo Fisher Scientific, Massachusetts, USA) according to the manufacturer’s instructions. The libraries were then sequenced on an Illumina NovaSeq instrument (Illumina) to generate 150-bp paired-end reads. The resulting raw reads were processed with Trimmomatic v0.39 to remove adapter sequences and to perform quality filtering. The processed reads were aligned to the Os-Nipponbare-Reference-IRGSP-1.0 reference genome sequence using BWA v0.7.18. The resulting SAM files were converted to BAM format, sorted, and indexed using SAMtools v1.9. SNP calling was performed using the mpileup and call commands in BCFtools v1.19. To restrict the analysis to previously characterized loci, a BED file containing the positions of 2,267,856 SNPs identified through parental analysis of the RIL population was provided. SNPs were called only at these specified positions and were used for all downstream analyses.

To remove redundant genotype information, SNPs were aggregated into haplotype blocks. Among the total 2,267,856 SNPs, SNPs were selected when at least one donor genome had nucleotides different from the common parent Hitomebore. By focusing on pairs of such SNPs (showing the coupling phase of the donor-type nucleotides), linkage disequilibrium (LD) was calculated using the RIL progeny showing segregation for the two SNPs. The SNPs that showed high LD (LD > 0.9) were then aggregated into one haplotype block. As a result, all SNPs were aggregated into 57,085 haplotype blocks. The detected SNPs in the breeding lines and their F2 progeny were also aggregated into the haplotype blocks. Principal component analysis (PCA) was performed to ascertain the genotypic diversity of the NAM population based on the genotypes of haplotype blocks.

### Genomic prediction

Multiple statistical models (GBLUP, Lasso, Elastic Net, Random Forest, Gradient boosting decision tree, BayesA, BayesB, and BayesC) were used to investigate optimal statistical models for accurate genomic prediction (*27–31*). The prediction accuracy of the genomic prediction model was calculated based on five-fold cross-validation using the NAM population. The Pearson’s correlation coefficient between the observed phenotypic values of the test dataset and the phenotypic values predicted from the genotyping data using the genomic prediction model was calculated and used to infer the prediction accuracy of the model. For five-fold cross-validation, all datasets were separated into training (80%) and test (20%) datasets so that each RIL population and phenotypic distribution were equally divided. GBLUP was performed using the R package “rrBLUP” v4.6.2 (*32*). Lasso, Elastic Net, and Random Forest were performed using the Python library “scikit-learn” v.1.2.0 (*33*). Gradient boosting decision tree was implemented using the Python library “LightGBM” v3.3.5 (*34*). BayesA, BayesB, and BayesC were implemented using the R package “BGLR” v1.1.4 (*35*). The hyperparameter optimization for Lasso, Elastic Net, Random Forest, and LightGBM was conducted by Bayesian optimization based on the training dataset using Optuna v3.2.0 (*36*). To investigate the changes in accuracy against the number of haplotype blocks provided during model construction, haplotype blocks were selected based on regression coefficients of Elastic Net. Genomic prediction models based on 3,000, 2,000, 1,000, 900, 800, 700, 600, 500, 400, 300, 200, 100, 75, 50, 25, and 10 selected haplotype blocks were built and compared for accuracy. For genomic prediction models used to predict other populations and to apply to genomic breeding, haplotype blocks were selected based on the mean values of regression coefficients that were derived from the Elastic Net models in all cross-validation folds. The prediction model that showed the highest accuracy in cross-validation was used to predict other populations. Genomic prediction models were built using 900 haplotype blocks for grain number, 1,000 haplotype blocks for panicle number, and 600 haplotype blocks for grain size (Data S2).

To confirm the robustness of the genomic prediction model for more complex genotypes derived from more than two parental lines, the phenotypic values of breeding lines for grain number, panicle number, grain size, and leaf width were predicted from the genotypes using the Elastic Net model. To validate breeding scheme, the phenotypic values of F2 individuals derived from RILs for grain number, panicle number, and grain size were predicted from the genotypes using the Elastic Net model.

To build genomic prediction models for crops other than rice, publicly available NAM datasets for maize (*Zea mays*), soybean (*Glycine max*), and sorghum (*Sorghum bicolor*) were used (*10*, *14*, *15*). SNP genotype and phenotype data were obtained from (https://www.panzea.org/data) for maize, from (https://soybase.org/data and https://www.soybase.org/SoyNAM/) for soybean, and from (doi:10.5061/dryad.gm073 and doi:10.5061/dryad.63h8fd4) for sorghum.

Representative agronomic traits with data covering at least two years and no obvious outliers were selected as the phenotype dataset. Ten traits were selected for maize: cob weight, days to tassel, ear diameter, ear row number, leaf length, leaf width, plant height, NIR oil (oil content determined by near-infrared spectroscopy), NIR protein, and NIR starch. The dataset of Aurora, Clayton, and Urbana were selected. Seven traits were selected for soybean: height, lodging, maturity, yield, protein, oil, and fiber. The dataset of Illinois, Indiana, and Nebraska were selected. Two traits were selected for sorghum: flowering time and plant height. The dataset of Manhattan was selected. Genomic prediction models using the Elastic Net model with haplotype blocks were built following the same pipeline as for the rice NAM population described above.

### Estimation of the genetic effect of genomic regions

The effects of genomic regions on grain number, panicle number, grain size, and tentative yield were calculated based on the regression coefficients of the Elastic Net model. To estimate the effect of genomic regions on the three traits, an Elastic Net model was built using 2,297 haplotype blocks merged from the haplotype blocks previously selected to build the most accurate model for each trait. To estimate genetic effects of genomic regions, five Elastic Net models were constructed from five training datasets used in a five-fold cross-validation. The mean values of the regression coefficients from five Elastic Net models were used to estimate effects. The effect of genomic regions on tentative yield was estimated by calculating Z-scores of the effects for haplotype blocks on grain number, panicle number, or grain size and adding up the Z-scores across all traits for each haplotype block (Data S3). The sum of regression coefficients or total Z-score values for haplotype blocks located every 1 Mb were aggregated as estimated genetic effects of genomic regions. Genomic regions that showed the top three positive effects on grain number and panicle number were identified as the candidate genomic regions of the donor genotype to be incorporated into an ideal target line from RILs (Data S4). Each genomic region was identified based on the estimated genetic effects with at most 4 Mb length, which is ∼1% of the whole rice genome, to retain a high proportion of genomic regions derived from the Hitomebore cultivar. The genomic regions that showed the top three negative effects on grain number, panicle number, or tentative yield were also identified as genomic regions where Hitomebore genotypes should be kept (Data S4).

### Selection of optimal RILs

The genetic effects of the genotypes from selected beneficial genomic regions or detrimental genomic regions were aggregated for each RIL and each trait (Data S4). The genetic effects on yield from selected detrimental genomic regions were also aggregated for each RIL (Data S4). RILs were selected as optimal RILs when they met the following criteria: (1) they showed the largest genetic effects from beneficial genomic regions among the RILs, (2) they had at least 65% of their genome derived from the Hitomebore cultivar (representing the top 10% of the NAM lines), and (3) they showed only small negative effects (if any) on yield from detrimental genomic regions (Data S4). For grain number, RILs N17-071 and N01-162, with the genotypes showing the largest positive aggregated genetic effect on grain number from three beneficial genomic regions, were selected (Data S4). For panicle number, RIL N08-011, with the genotype showing the largest positive aggregated genetic effect on panicle number from two beneficial genomic regions, was selected (Data S4). Combinations of N01-162 and N08-011 or N17-071 and N08-011 were used as starting materials in breeding schemes, to limit the number of that as few donor genotypes to be introduced as possible to obtain large positive genetic effects.

### Simulation of breeding plan

Recombination events were simulated based on the observed recombination numbers across the NAM population. The number of recombination events on each chromosome was simulated based on the Poisson distribution. The parameter lambda of the Poisson distribution was set to match the distribution of the number of recombination events in the simulated RIL population and that of the observed number of recombination events in the NAM population. The match of distributions of the number of recombination events between the observed and predicted populations were tested by a Kolmogorov–Smirnov (K-S) test for goodness of fit. *K*/2, where *K* is the number of recombination events for each chromosome in the NAM population, was used as parameter lambda, since the simulated distributions did not differ from the observed distributions based on the K-S test (*p* > 0.05) in the half of the 12 chromosomes (Fig. S11). The positions of recombination events were simulated based on the probability density function estimated using Gaussian kernels from the positions of observed recombination events in the NAM population (Fig. S11).

The generation of an F2 progeny from crosses derived from selected RILs and Hitomebore, crossing between F2 individuals, and self-pollination in the breeding plan were simulated. In the simulation, markers were assumed to be located in ∼2.5-Mb intervals in each genomic region derived from the donor cultivars to select lines for the next crossing stage (Fig. S12). We also assumed that 100 plants were generated per cross; the recombination landscape was simulated as described above. The number of progeny with donor genotypes of all beneficial genomic regions was counted based on marker genotypes (Fig. S12). Among the progeny plants with donor genotypes for markers at all beneficial genomic regions, a plant with the largest representation of the Hitomebore genome, as judged by the genotypes of other markers, was advanced to the next stage for crossing. A simulation of the entire breeding plan was performed 100 times. The genotypes of the target cultivars generated by 100 simulated breeding plans were provided to the genomic prediction model to predict the phenotype values. The mean values and standard deviation values of the predicted phenotype values were calculated and compared to the predicted phenotype values of Hitomebore (Fig. S10).

## Acknowledgments

We thank Yumiko Ogasawara, Hiroyuki Kanzaki, Eiko Kanzaki, and all others who supported this study. We also thank the National Agriculture and Food Research Organization (NARO) Genebank, Japan, for providing rice accessions. Part of the analysis was performed using the National Institute of Genetics (NIG) supercomputer at the Research Organization of Information and Systems (ROIS) National Institute of Genetics and the Academic Center for Computing and Media Studies (ACCMS) supercomputer at Kyoto University.

## Funding

The Research Program on the Development of Innovative Technology from the Project of the Bio-oriented Technology Research Advancement Institution (BRAIN) (JPJ007097)

JSPS KAKENHI (20H02962)

JSPS KAKENHI (23H02189)

JSPS KAKENHI (JP23K26882)

## Author contributions

Examples:

Conceptualization: T.S., A.A., R.T.

Methodology: T.S., A.A.

Investigation: T.S., A.A., H.T., H.Y.,

S.Natsume., K.O., H.U., K.I., T.F., Y.O.,

S.Nakajo., M.S., T.T.

Formal analysis: T.S.

Data curation: T.S., A.A.

Funding acquisition: A.A., R.T.

Supervision: A.A., R.T.

Writing – original draft: T.S.

Writing – review & editing: T.S., A.A., R.T.

## Data, code, and materials availability

The genotype and phenotype data used in this study were deposited to the Zenodo repository (https://doi.org/10.5281/zenodo.15835034). All sequence data used in this study were deposited under accession number PRJDB13864. All scripts used in this study were deposited to the GitHub repository (https://github.com/slt666666/genomic_prediction_scripts).

