## Supplemental Materials for "Genomic prediction models based on a large-scale recombinant population allow rapid breeding of desired genotypes"

Toshiyuki Sakai *et al.*

**This PDF file includes:**

Figs. S1 to S14
Table S1

**Other Supplementary Materials for this manuscript include the following:**

Data S1 to S4

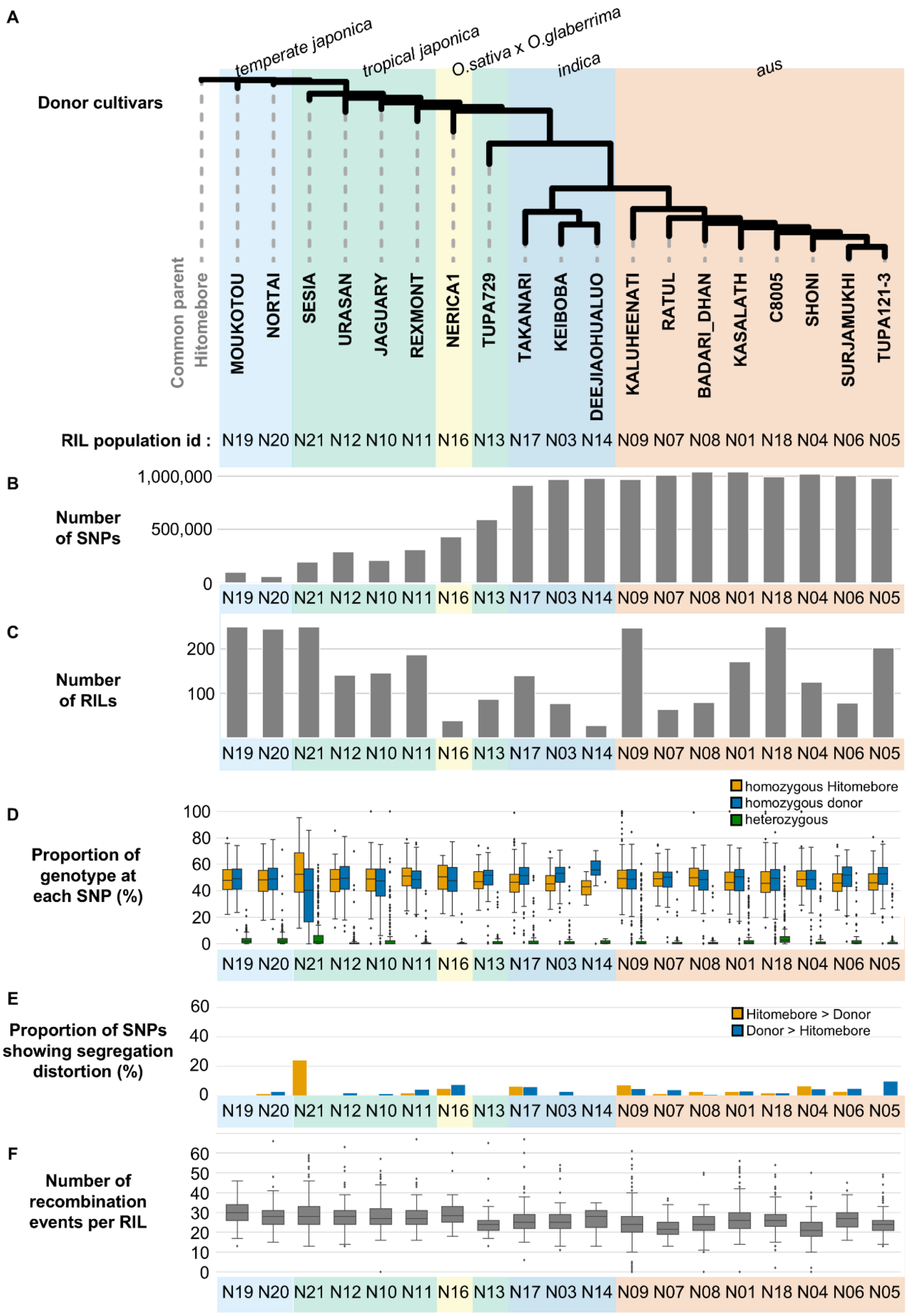


**Fig. S1. Summary of the NAM population.** (**A**) Unrooted phylogenetic tree, reconstructed based on single-nucleotide polymorphisms (SNPs) between the common parental line and the donor cultivars. (**B**) Number of SNPs between the common parent (Hitomebore) and each donor cultivar. (**C**) Number of recombinant inbred lines (RILs) derived from each cross. (**D**) Proportion of the three types of genotypes (homozygous for Hitomebore, heterozygous, homozygous for the donor parent) in each RIL. (**E**) Proportion of SNPs showing segregation distortion in each RIL population. (**F**) Number of recombination events in each RIL population.

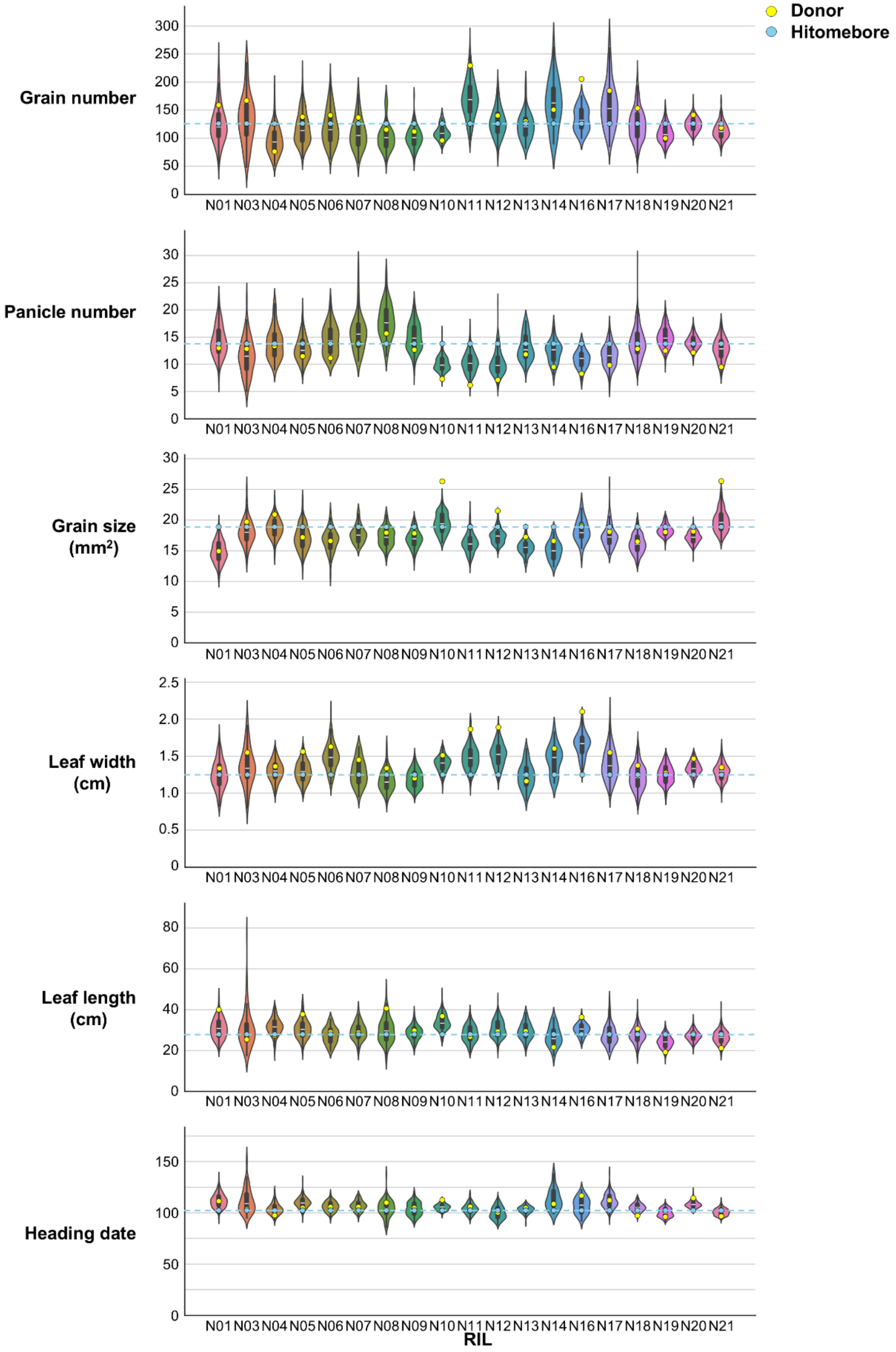
Donor
Hitomebore
Grain number
Panicle number
Grain size (mm²)
Leaf width (cm)
Leaf length (cm)
Heading date
N01 N03 N04 N05 N06 N07 N08 N09 N10 N11 N12 N13 N14 N16 N17 N18 N19 N20 N21
RIL

**Fig. S2. Distribution of phenotypic values for each RIL population.** Violin plots of the phenotypic values for each trait and each RIL population. Yellow and blue dots indicate the phenotypic values for the donor cultivar and Hitomebore, respectively.

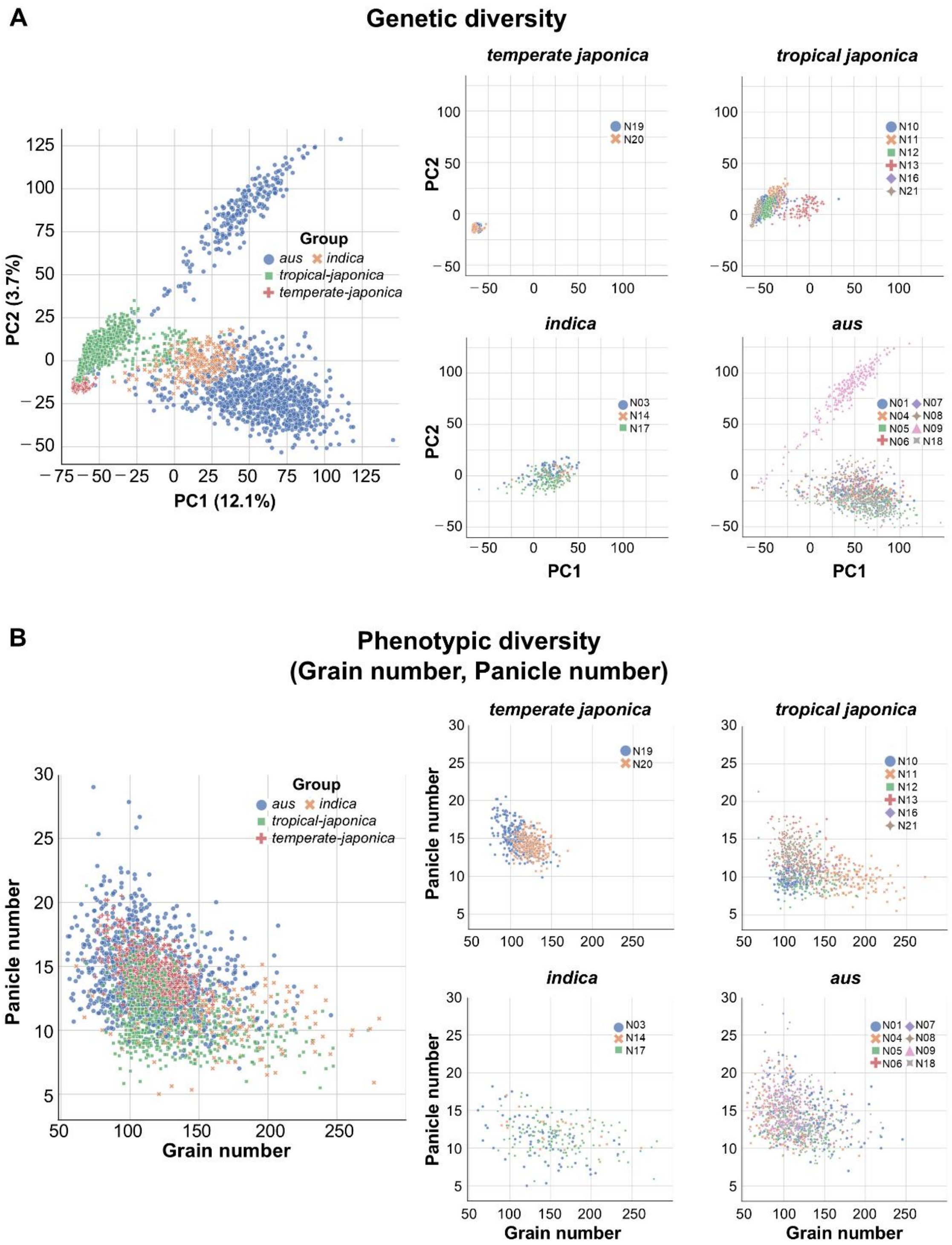


**Fig. S3. Genetic and phenotypic diversity of the NAM population.** (**A**) Genetic diversity of the entire rice NAM population (left) or subpopulations (right: *temperate japonica*, *tropical japonica*, *indica*, *aus*), as estimated by principal component analysis (PCA) based on the genotypes at all haplotype blocks across the NAM population. (**B**) Scatterplot showing the phenotypic diversity for grain number and panicle number among the entire rice NAM population (left) or subpopulations (right: *temperate japonica*, *tropical japonica*, *indica*, *aus*).

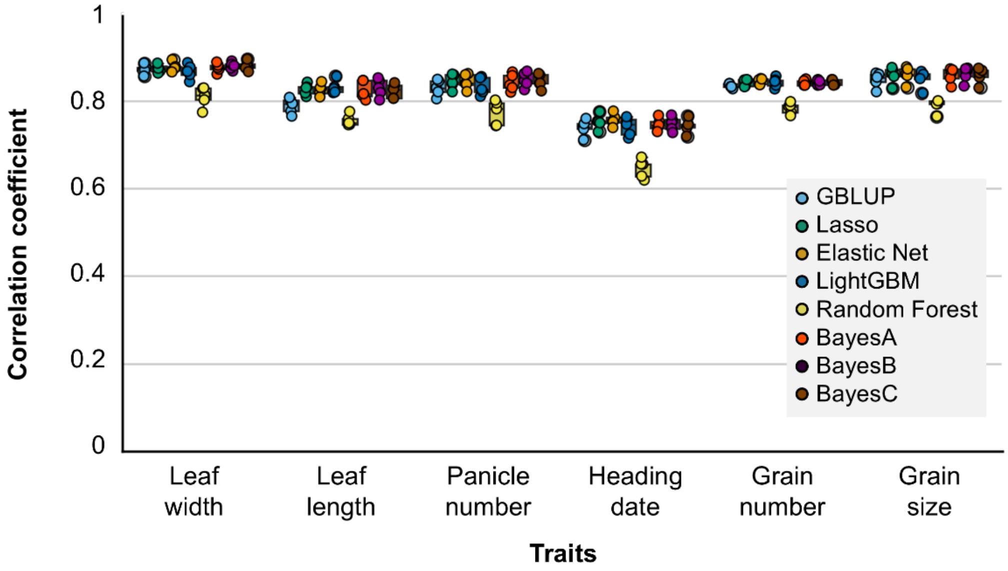


**Fig. S4. Accuracy of genomic prediction from each of eight statistical models.** Boxplots showing the prediction accuracy of five-fold cross-validation for each trait. The *x*-axis shows each trait; the *y*-axis shows the Pearson's correlation coefficient between the predicted and measured phenotypic values of the test dataset. Each statistical model is indicated in a different color.

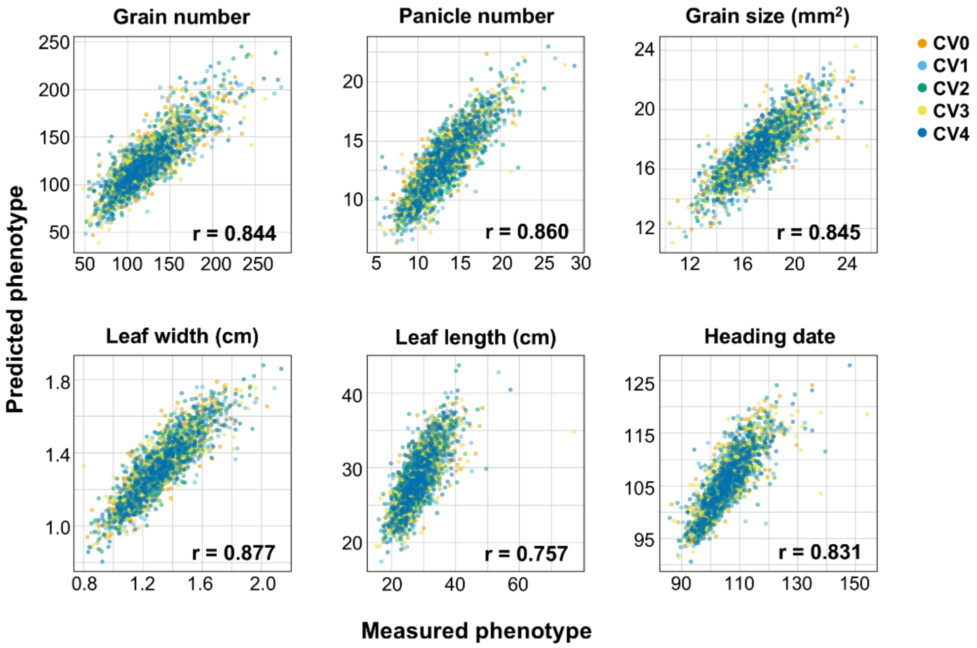


**Fig. S5. Relationships between measured phenotypic values and predicted values obtained by the Elastic Net model.** This figure shows the data presented in Figure 3B with the addition of the results of leaf length and heading date. Scatterplots showing the relationship between measured and predicted phenotypic values of five-fold cross-validation for each trait. The *x*-axis shows the measured phenotypic values; the *y*-axis shows the predicted phenotypic values. The average Pearson's correlation coefficients across the five cross-validations (CVs) are given at the bottom right of each plot.

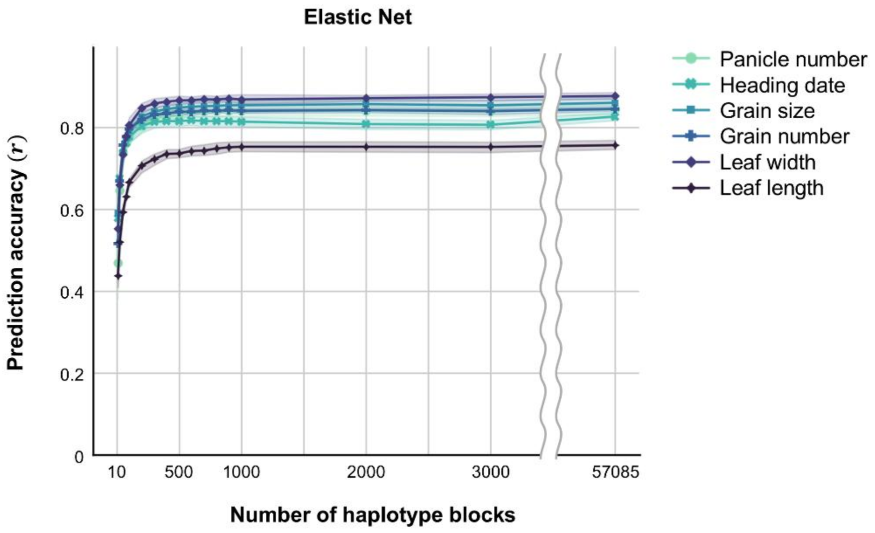


**Fig. S6. Estimation of prediction accuracy by genomic prediction models depending on the number of haplotype blocks.** Prediction accuracy of Elastic Net models as a function of the number of haplotype blocks. The *x*-axis shows the number of haplotype blocks included in each model; the *y*-axis shows the Pearson's correlation coefficient between predicted and measured phenotypic values of the test dataset.

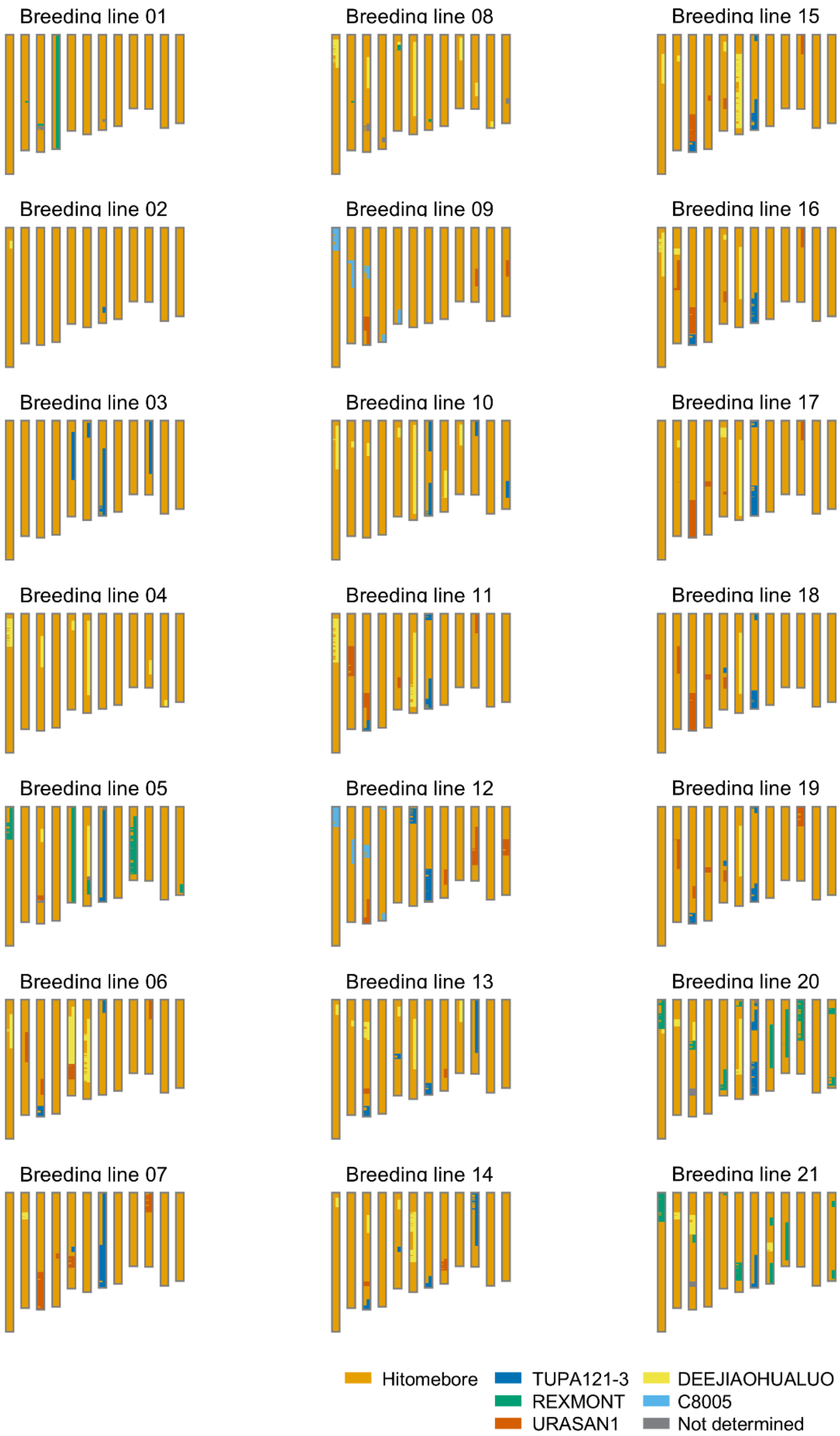


**Fig. S7. Graphical representation of the genotypes from the 21 rice lines used for validation of the accuracy of models applied to lines with more than two donor genomes.** Each panel shows the graphical representation of the genotype for the indicated breeding line. The genomic regions derived from Hitomebore are shown in orange. The genomic regions derived from donor cultivars are shown in a different color, as indicated at the bottom of the figure.

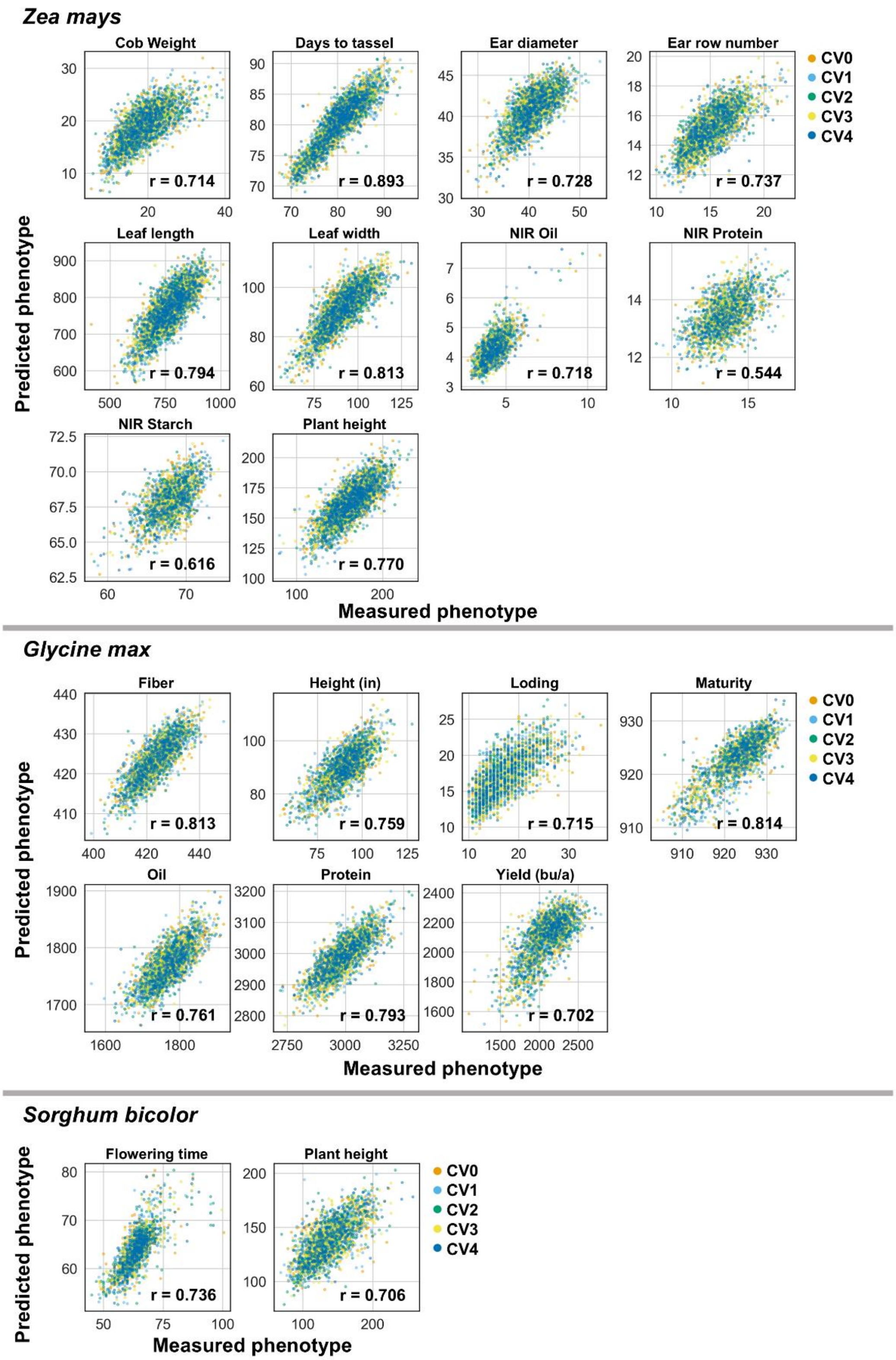


**Fig. S8. Prediction accuracy of the models applied to NAM populations for maize, soybean, and sorghum.** Scatterplots showing the relationship between measured and predicted phenotype values for the indicated phenotypes for the NAM population of maize (top), soybean (middle), and sorghum (bottom). The average Pearson's correlation coefficients across the five cross-validations (CVs) are given at the bottom right of each plot.

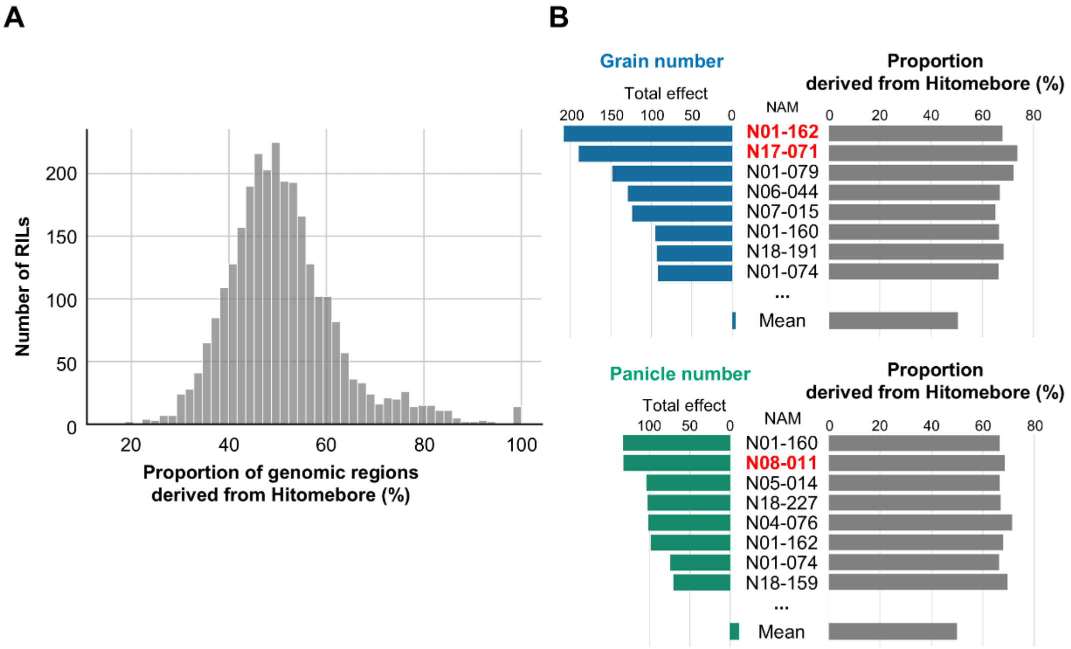


**Fig. S9. Proportion of genomic regions derived from Hitomebore among the RILs.** (**A**) Distribution of the proportion of genomic regions derived from Hitomebore for 2,787 RILs. (**B**) Bar graphs showing the total genetic effect on a target trait (left) and the proportion of genomic regions from Hitomebore (right), for the top eight RILs with the highest total positive genetic effect on the target trait.

### Figures

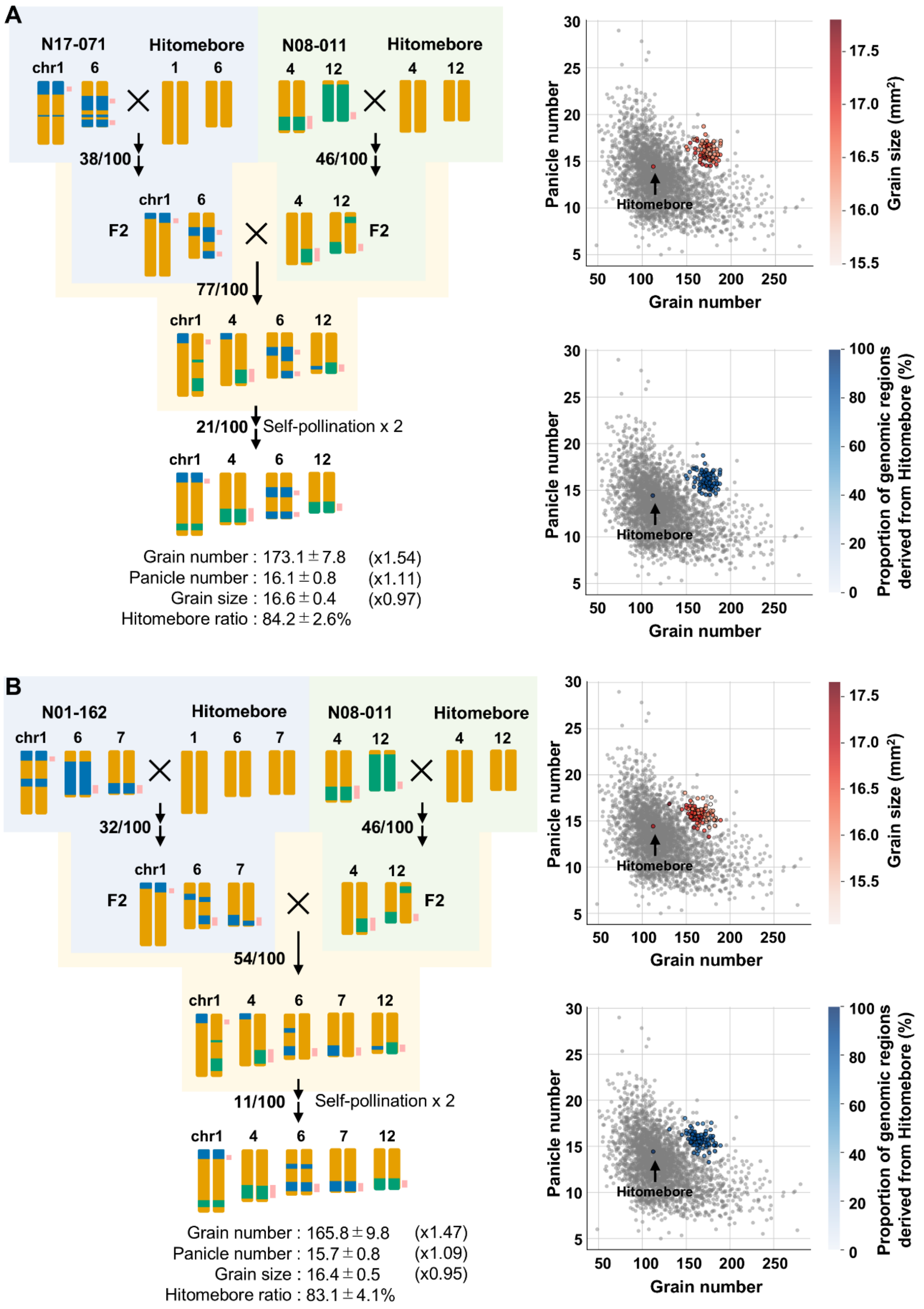
A
N17-071
Hitomebore
N08-011
Hitomebore
chr1
6
1
4
12
38/100
46/100
F2
77/100
21/100
Self-pollination x 2
Grain number : 173.1 ± 7.8 (x1.54)
Panicle number : 16.1 ± 0.8 (x1.11)
Grain size : 16.6 ± 0.4 (x0.97)
Hitomebore ratio : 84.2 ± 2.6%
B
N01-162
7
32/100
54/100
11/100
Grain number : 165.8 ± 9.8 (x1.47)
Panicle number : 15.7 ± 0.8 (x1.09)
Grain size : 16.4 ± 0.5 (x0.95)
Hitomebore ratio : 83.1 ± 4.1%
Panicle number
Grain number
Hitomebore
Grain size (mm²)
Proportion of genomic regions derived from Hitomebore (%)

**Fig. S10. Simulation results of the breeding plans.** (**A**) Simulation results of a breeding plan using N17-071 and N08-011 as parents. Each RIL was crossed to Hitomebore; F2 plants derived from each cross are then crossed, followed by two rounds of self-pollination. This figure shows the data presented in Figure 4C and D with the additional results of the simulation. (**B**) Simulation results of a breeding plan using N01-162 and N08-011 as parents. In the diagrams of the breeding plan, the number of progeny that carry donor genotypes at all beneficial genomic regions among 100 simulated progeny (XX/100) are given to the left of the arrows after each cross. The average and standard deviation of the predicted phenotypic values of the target cultivars produced by the breeding plan are shown below the breeding scheme, as well as the fold-increase from the predicted phenotypic value of Hitomebore. The top right scatterplots of (A) and (B) show the distribution of 100 simulated phenotypic values for grain number, panicle number, and grain size. The bottom right scatterplots of (A) and (B) show the distribution of the proportion of genomic regions derived from Hitomebore as colored dots. Gray circles show measured phenotypic values for the RILs in the NAM population.

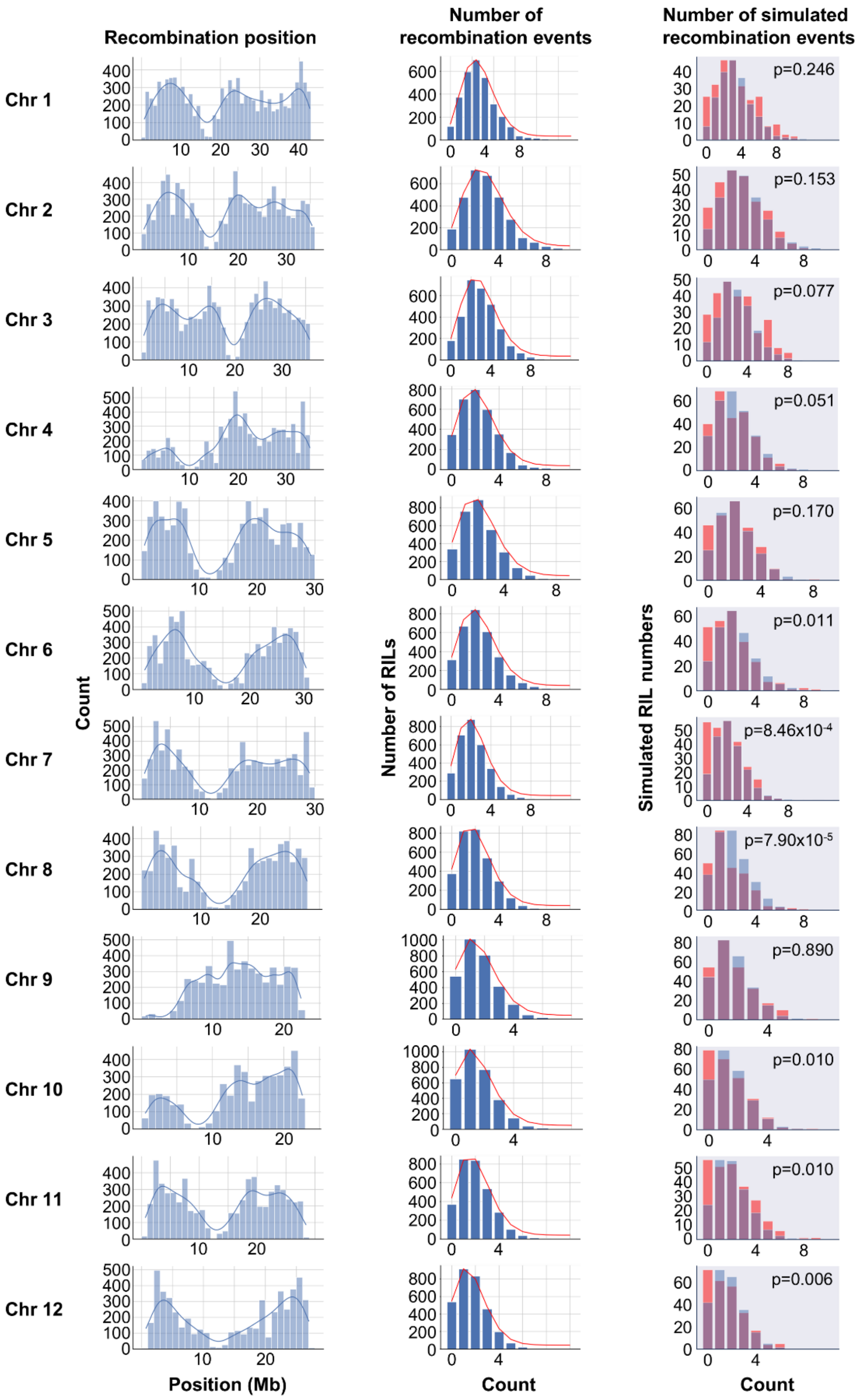


**Fig. S11. Observed recombination events in the NAM population and simulated recombination events.** Left, positions of observed recombination events on each chromosome. Middle and right, number of observed (middle) or simulated (right) recombination events. Red lines show the approximate Poisson distributions (middle). Blue bars, observed recombination events; red bars, simulated recombination events (right). The p-values for each Kolmogorov–Smirnov (K-S) test are given on the right. The number and positions of recombination events per cross were simulated based on these probability distributions. The genotype of each plant was determined based on the genotype of the parental lines and simulated positions and numbers of recombination events.

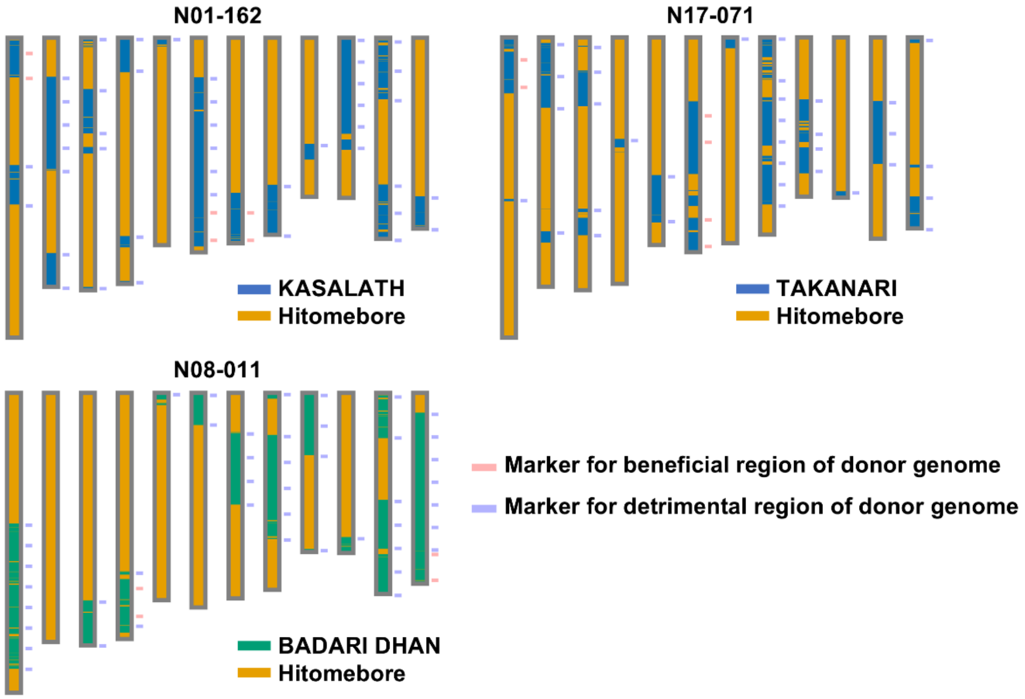


**Fig. S12. Position of DNA markers designed to perform rapid breeding.** Graphical representation of the genotypes for the lines N01-162, N08-011, and N17-071. Blue and red lines show the positions of markers. In the simulation of the breeding plan, plants with donor genotypes at all markers in beneficial regions are first selected. Among these selected plants, those with the highest number of markers with the Hitomebore genotype in detrimental regions are selected for the next cross.

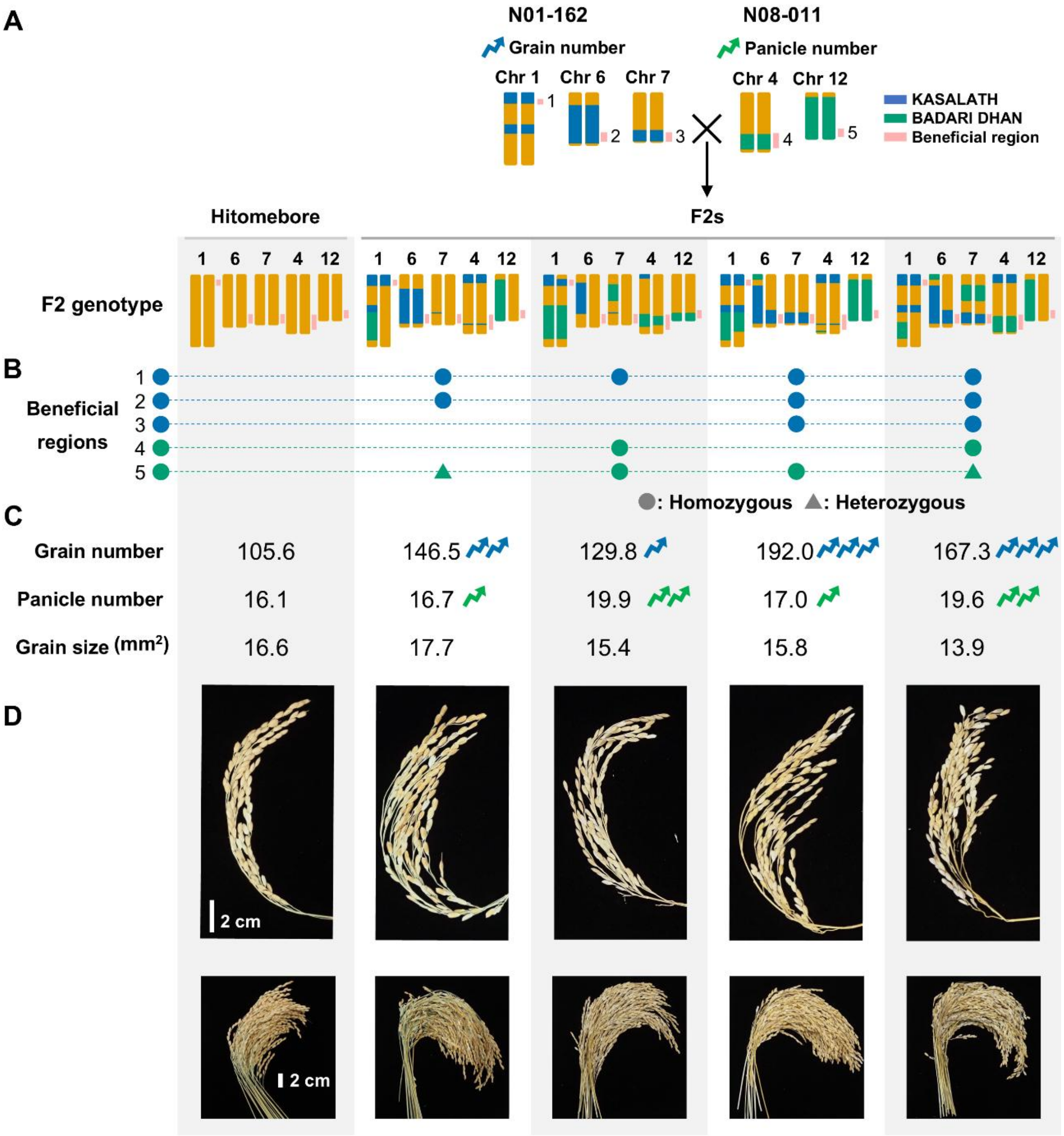


**Fig. S13. Phenotypic values of F2 plants derived from a cross between N01-162 and N08-011.** (**A**) Top, diagram of the cross and generation of the F2 population. (**B**) Graphical representation of the genotypes at each target beneficial genomic region. (**C**) Phenotypic values of grain number, panicle number, and grain size. (**D**) Representative photographs of panicles from F2 plants with each genotype.

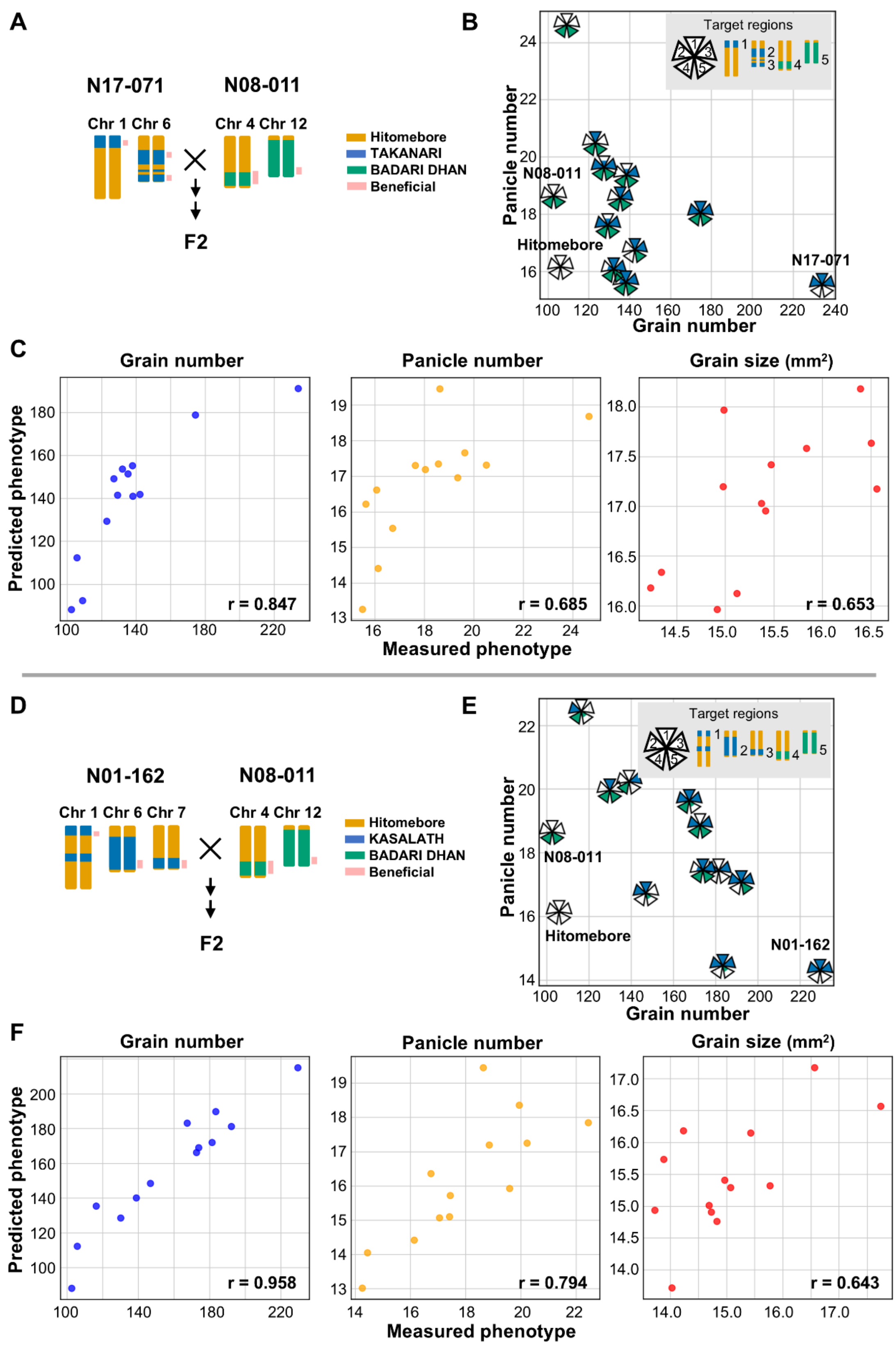


**Fig. S14. Phenotypic values and prediction accuracy for F2 genotypes derived from crosses between selected RILs.** (**A, D**) Diagrams of the cross and generation of the F2 population. (**B,**

**E**) Scatterplots of the phenotypic values for the parental lines and specific groups of F2 plants with the indicated genotypes for grain number and panicle number. (**C, F**) Scatterplots showing the relationship between measured and predicted phenotypic values for F2 plants, parental lines, and Hitomebore. (**A–C**) F2 plants derived from a cross between N17-071 and N08-011. (**D–F**) F2 plants derived from a cross between N01-162 and N08-011.

**Table S1.** Donor cultivar combinations used for the development of breeding lines for genomic prediction validation.

| ID | Donor varieties |
|---|---|
| Breeding line 01 | TOYAMA 73 |
| Breeding line 02 | DUNGHAN SHALI |
| Breeding line 03 | TUPA 121-3 |
| Breeding line 04 | DEEJIAOHUALUO |
| Breeding line 05 | TOYAMA 73, REXMONT, TUPA 121-3, DEEJIAOHUALUO |
| Breeding line 06 | TUPA 121-3, DEEJIAOHUALUO, URASAN 1, DUNGHAN SHALI |
| Breeding line 07 | TUPA 121-3, DEEJIAOHUALUO, URASAN 1, DUNGHAN SHALI |
| Breeding line 08 | DEEJIAOHUALUO, TOYAMA 73 |
| Breeding line 09 | URASAN 1, DUNGHAN SHALI, C8005, TUPA 121-3 |
| Breeding line 10 | DEEJIAOHUALUO, TUPA 121-3 |
| Breeding line 11 | DEEJIAOHUALUO, TUPA 121-3, URASAN 1, DUNGHAN SHALI |
| Breeding line 12 | URASAN 1, DUNGHAN SHALI, C8005, TUPA 121-3 |
| Breeding line 13 | URASAN 1, DUNGHAN SHALI, DEEJIAOHUALUO, TUPA 121-3 |
| Breeding line 14 | URASAN 1, DUNGHAN SHALI, DEEJIAOHUALUO, TUPA 121-3 |
| Breeding line 15 | DEEJIAOHUALUO, TUPA 121-3, URASAN 1, DUNGHAN SHALI |
| Breeding line 16 | DEEJIAOHUALUO, TUPA 121-3, URASAN 1, DUNGHAN SHALI |
| Breeding line 17 | DEEJIAOHUALUO, TUPA 121-3, URASAN 1, DUNGHAN SHALI |
| Breeding line 18 | DEEJIAOHUALUO, TUPA 121-3, URASAN 1, DUNGHAN SHALI |
| Breeding line 19 | DEEJIAOHUALUO, TUPA 121-3, URASAN 1, DUNGHAN SHALI |
| Breeding line 20 | TOYAMA 73, REXMONT, TUPA 121-3, DEEJIAOHUALUO |
| Breeding line 21 | TOYAMA 73, REXMONT, TUPA 121-3, DEEJIAOHUALUO |

**Data S1. (separate file).** This file contains the tabulated data of characteristics of RIL populations in the NAM population behind the figure S1.

**Data S2. (separate file).** This file contains the tabulated data of the prediction accuracy of genomic prediction models based on the NAM population behind the figures 3, S4, S5, S6.

**Data S3. (separate file).** This file contains the tabulated data of the estimated effect on grain number, panicle number, grain size, total yield of important haplotype blocks behind the figure 4A.

**Data S4. (separate file).** This file contains the tabulated data of the estimated effect of selected genomic regions for each RIL behind the figures 4B, S9.
